# Insights into chloroplast genome variation across Opuntioideae (Cactaceae)

**DOI:** 10.1101/2020.03.06.981183

**Authors:** Matias Köhler, Marcelo Reginato, Tatiana T. Souza-Chies, Lucas C. Majure

## Abstract

Chloroplast genomes (plastomes) are frequently treated as highly conserved among land plants. However, many lineages of vascular plants have experienced extensive structural rearrangements, including inversions and modifications to the size and content of genes. Cacti are one of these lineages, containing the smallest plastome known for an obligately photosynthetic angiosperm, including the loss of one copy of the inverted repeat (∼25 kb) and the *ndh* genes suite, but only a few cacti from the subfamily Cactoideae have been sufficiently characterized. Here, we investigated the variation of plastome sequences across the second-major lineage of the Cactaceae, the subfamily Opuntioideae, to address 1) how variable is the content and arrangement of chloroplast genome sequences across the subfamily, and 2) how phylogenetically informative are the plastome sequences for resolving major relationships among the clades of Opuntioideae. Our *de novo* assembly of the *Opuntia quimilo* plastome recovered an organelle of 150,347 bp in length with both copies of the inverted repeats and the presence of all the *ndh* genes suite. An expansion of the large single copy unit and a reduction of the small single copy was observed, including translocations and inversion of genes as well as the putative pseudogenization of numerous loci. Comparative analyses among all clades within Opuntioideae suggested that plastome structure and content vary across taxa of this subfamily, with putative independent losses of the *ndh* gene suite and pseudogenization of genes across disparate lineages, further demonstrating the dynamic nature of plastomes in Cactaceae. Our plastome dataset was robust in determining relationships among major clades and subclades within Opuntioideae, resolving three tribes with high support: Cylindropuntieae, Tephrocacteae and Opuntieae. A plastome-wide survey for highly informative phylogenetic markers revealed previously unused regions for future use in Sanger-based studies, presenting a valuable dataset with primers designed for continued evolutionary studies across Cactaceae. These results bring new insights into the evolution of plastomes in cacti, suggesting that further analyses should be carried out to address how ecological drivers, physiological constraints and morphological traits of cacti may be related with the common rearrangements in plastomes that have been reported across the family.

## 1. Introduction

Cacti comprise one of the most charismatic plant clades of the world, exhibiting an enormous variety of growth forms, morphology and intriguing niche occupancy across the Americas (Britton & Rose 1919; Anderson 2001; Hunt et al. 2006; Hernández-Hernández et al. 2011). Members of the family are conspicuous elements of the arid and semiarid succulent biome of the New World, also inhabiting subtropical and tropical forests (Taylor and Zappi 2004; Hunt et al. 2006). This distributional pattern is accompanied by high species diversity with heterogeneous diversification rates across evolutionary lineages (Arakaki et al. 2011; Hernández-Hernández et al. 2014).

Some uncommon features in most Angiosperms, such as succulent tissues, Crassulacean acid metabolism (CAM), betalain pigments and the reduction of or absence of leaves are typical characters of cacti that have long captured the attention of plant biologists and have been suggested as adaptations to allow survival in harsh environments (Mooney et al. 1977; Mauseth 1999; Landrum 2002; Nobel 2002). Besides major morphological and physiological adaptations, genetic and genomic-level changes derived from host of selective pressures are also expected to be present. For example, whole genome duplication events have long been suggested to be associated with adaptations to extreme environments (e.g., Stebbins 1971; Soltis & Soltis 2000; Brochmann et al. 2004), and significant gene family expansion in genes related to stress adaptation, as well as more restricted events of gene duplications were reported in lineages of Caryophyllales adapted to severe environments including in cacti (Wang et al. 2019).

Although gene content, structural organization and size of the chloroplast genome (plastomes) of land plants is often considered highly conserved (Raubeson & Jansen 2005; Chumley et al. 2006; Wicke et al. 2011), deviations have been increasingly reported in some clades and have challenged the generality of this phenomenon (Daniell et al 2016; Ruhlman and Jansen 2018; Mower and Vickrey 2018). Astonishing variety of size have been observed across land plants, from 19 kb in a non-photosynthetic *Epipogium roseum* (Orchidaceae) to giant plastomes with 217 kb, as in *Pelargonium × hortorum* (Geraniaceae) (Chumley et al 2006; Schelkunov et al 2015), reflected by expansions or contraction of the inverted repeat (IR), large single copy (LSC) or even small single copy (SSC) units. Also, the independent losses of one copy of the inverted repeat region (∼25 kb in size) have been identified across disparate clades, such as Fabaceae, Geraniaceae, Orobanchaceae and Cactaceae (Cai et al. 2008; Ruhlman & Jansen 2014; Sanderson et al. 2015), and a variety of taxa have lost particular genes in other parts of the genome. For instance, most or all of the suite of 11 functionally related *ndh* genes were lost in some parasites, carnivorous plants and xerophytes (Braukmann et al., 2009; Wicke et al., 2011; Iles et al., 2013; Peredo et al., 2013; Ruhlman et al., 2015; Sanderson et al. 2015).

Members of Cactaceae also have experienced different alterations in their chloroplast genome. A conserved inversion of ∼6 kb on the large single copy unit comprising the *trnM*-*rbcL* genes have long been suggested (Wallace 1995) and more recently confirmed (Sanderson et al. 2015; Solorzano et al. 2019; Majure et al. 2019). Besides that, the first cactus plastome assembled from the saguaro cactus (*Carnegiea gigantea* (Engelm.) Britton & Rose) exhibited an exceptional reduction in size (113 kb) and gene content, including the loss of one of the two inverted repeat regions and nine of the 11 *ndh* genes (Sanderson et al. 2015). More recently, newly assembled plastomes of seven species of the short-globose cacti of *Mammillaria* revealed three different plastome structures across the genus, all with two copies of a divergent inverted repeat, including (i) an extreme reduction of IRs (< 1 kb); (ii) an intermediate reduction of IR with translocation of some typical LSC genes to the IR; and (iii) a structure with a divergent IR structure and a surprisingly reduced plastome (∼107 kb), being now the putative smallest plastome known for an obligately photosynthetic angiosperm (Solorzano et al. 2019). Majure et al. (2019), assembling chloroplast genomes for phylogenetic studies, found that *ycf1* and *ycf2* were pseudogenized, and that *ycf1* gene had been translocated adjacent to the *rpl32* gene in *Cylindropuntia bigelovii* (Engelm.) F.M.Knuth (Opuntioideae).

The taxonomy and systematics of Cactaceae has a long history of Linnean hierarchical classification based on morphological characters (Schumann 1899; Britton and Rose 1919; Backeberg 1958; Hunt et al. 2006). However, only with the advance of modern techniques adopting molecular data to reconstruct phylogenies, have systematic biologists been able to test and build more reliable hypothesis about the relationships of this peculiar group, which is replete with homoplasy (Hernández-Hernández et al. 2011). Three major well supported clades are currently circumscribed as subfamilies: Opuntioideae, Maihuenioideae and Cactoideae; while the traditional “Pereskioideae” subfamily has been revealed as a basal grade including the two leafy lineages of the cacti, which are subsequent sisters to the rest, i.e. *Leuenbergia* and *Pereskia* (Edwards et al. 2005, reviewed in Guerrero et al. 2018).

Opuntioideae (∼350 spp.) is the most widespread subfamily with members occurring from southern South America (Patagonia) to northern North America (Saskatchewan, Canada) (Britton & Rose 1919; Anderson 2001; Hunt et al. 2006; Ritz et al. 2012; Majure & Puente 2014; Majure et al. 2019). The group shows interesting morphological synapomorphies, such as the small brushlike, barbed spines (i.e., glochids) and the bony aril surrounding a campylotropous ovule (Taylor et al. 2002; Stuppy 2002). However, members of Opuntioideae have a long blurred taxonomic history with different approaches applied to circumscribe and to delimit taxa across different taxonomic levels, from species to tribes (Schumann 1899; Britton & Rose 1919; Taylor & Stuppy 2002; Stuppy 2002; Hunt 2002). Traditional classifications based on general morphology have divided the subfamily from one genus (*Opuntia* s.l. (L.) Mill.) to up to 20 smaller genera (Britton and Rose 1919; Hunt et al. 2006). Nonetheless, molecular phylogenetic studies showed that *Opuntia* s.l was paraphyletic, which led to the recognition of numerous smaller genera corresponding to well-supported clades (Taylor and Stuppy 2002). Likewise, the tribal classification of Opuntioideae has been controversial based on different approaches, with up to six tribes proposed (Doweld, 1999; Wallace and Dickie, 2002; Hunt 2011). Although previous studies have improved our understanding of the relationships among the major clades in Opuntioideae (Griffith & Porter 2009; Ritz et al. 2012; Majure et al. 2019), support for the relationships among those clades still needs to be strengthened.

Apart from the external and internal transcribed spacer (ETS and ITS) of the nuclear ribosomal repeats (NRR) and *ppc* marker, most molecular phylogenies of cacti have been historically based on few plastid markers *(trnL-trnF*, *rpl16*, *trnK* and *matK*) (Nyffeler 2002; Edwards et al. 2005; Korotova et al. 2010; Demaio et al. 2011; Arakaki et al. 2011; Bárcenas et al. 2011; Hernández-Hernández et al. 2011; Hernández-Hernández et al. 2014; Ritz et al. 2012; Bárcenas 2016; Luna-Vargas et al. 2018). While these markers have shown to be potentially able to resolve some clades, some relationships are still lacking support (Nyffeler 2002; Griffiths & Porter 2009; Hernández-Hernández et al. 2011; Bárcenas et al. 2011). In this case, next-generation sequencing (NGS) could be a useful tool, since it has transformed the study of nonmodel plant taxa in phylogenetic inferences with high throughput data allowing deep resolutions across major plant clades (Xi et al. 2012; Ma et al. 2014; Gardner et al. 2016; Zong et al. 2019). NGS data are also showing to be extremely useful for discovering informative regions across genomes, for marker development (Wu et al. 2010; Dong et al. 2012; Ripma et al. 2014; Reginato et al. 2016), as well as to investigate chloroplast genome evolution (Dong et al. 2013; Mower & Vickrey 2018; Yao et al. 2019). Nevertheless, this approach is still in its infancy across Cactaceae (Majure et al. 2019) and remains a path to be explored.

Here, we investigate the use of next-generation sequencing across Opuntioideae to address two major questions: (1) how homogenous is the content and arrangement of chloroplast genomes across the subfamily? and (2) how phylogenetically informative are chloroplast genome sequences for resolving major relationships among the clades of Opuntioideae? We used a combination of *de novo* and reference-guided assemblies to process genome skimming data: (i) assembling and characterizing the first chloroplast genome of an *Opuntia* species, *O. quimilo,* (ii) investigating overall patterns of reference-guided assemblies and comparative chloroplast genome sequence analyses across the subfamily, (iii) inferring phylogenetic relationships with assembled sequences and (iv) surveying plastomes for highly informative phylogenetic markers for Sanger-based studies for future use.

## 2. Material and methods

### 2.1 Taxon sampling, DNA extraction and sequencing

All currently recognized genera in Opuntioideae (sensu Hunt et al. 2006, plus Majure et al. 2019 for *Grusonia* s.l.) were sampled with one accession per genus, resulting in a dataset of 17 taxa which were sequenced via genome-skimming (Straub et al., 2012; Majure et al. 2019). Three additional samples were selected as outgroup taxa (Cactoideae: *Parodia magnifica* and *Coryphantha macromeris*; and *Pereskia*: *Pereskia sacharosa*). Plant materials were from wild collections or from the Desert Botanical Garden’s living collection (see Table S1 for details). DNA was extracted from silica-dried or fresh epidermal tissues using a standard CTAB incubation (Doyle & Doyle 1987) followed by chloroform/isoamyl alcohol precipitation and silica column-based purification steps, as described in Neubig et al. (2014) and Majure et al. (2019). Whole genomic DNAs were quantified using the Qubit dsDNA BR Assay Kit and Qubit 2.0 Fluorometer (Life Technologies, Carlsbad, California, USA); high-molecular-weight DNA (>15 kb) samples showing no degradation were considered suitable and sent to Rapid Genomics LLC (http://rapidgenomics.com/home/; Gainesville, FL.) for library preparation and high-throughput sequencing using the Illumina HiSeq X platform with 150 bp paired-end reads.

### 2.2. De novo assembly and data processing for chloroplast genome sequences

Raw reads were imported into Geneious 11.1.5 (Biomatters, Auckland, New Zealand), and paired reads were set with an expected insert size of 300 bp calculated with BBMap (Bushnell 2016). Low quality bases (Q < 20) were trimmed and all reads shorter than 20 bp were discarded using BBDuk for quality control (Bushnell 2016). Different methods were employed to assemble the chloroplast genome of the diploid *Opuntia quimilo* K. Schum. First, a *de novo* assembly was performed with 40% of the reads using the Geneious *de novo* assembler (low/fast sensitivity option). A consensus sequence (with a majority threshold for sequence matching - fewest ambiguities) of each contig greater than 1,000 bp in length was saved. Considering that the Cactaceae plastomes already published have unusual rearrangements, we looked for plastid contigs searching those saved contigs against the *Portulaca oleracea* L. plastome (Portulacaceae, one of the closest relatives of Cactaceae; see Walker et al. 2018) (GenBank accession KY490694, Liu et al. 2017) using MegaBLAST (adapted from Ripma et al.2014). Additional chloroplast genome *de novo* assemblies were performed using a set of different pipelines, such as GetOrganelle (Jin et al. 2019) and NOVOPlasty (Dierckxsens et al. 2017) to cross-validate and compare the assemblies. After checking the convergence of the assemblies from the different pipelines and the plastid contig recovered from the Geneious *de novo* assembly, we used the NOVOPlasty circular contig for downstream analyses. Annotations were performed with GeSeq (Tillich et al. 2017), using default parameters to predict protein-coding genes by HMMER profile search and ARAGORN v1.2.38 (Laslett and Canback, 2004); tRNA genes were annotated with tRNAscan-SE v2.0 (Lowe and Eddy, 1997), and BLAST searches were used to annotate ribosomal RNA (rRNA), tRNA and DNA genes conserved in embryophyte plastomes (Wommack et al., 2008). All annotations were cross checked with the “Annotate from” feature in Geneious, transferring annotations with a 50% or greater similarity from the *P. oleracea* plastome, and eventual start/stop codons were manually adjusted with the “Open Read Frame (ORF)” feature from Geneious. The genes that had their structures affected by the insertion of many internal stop codons or ORF, thus did not forming their respective full coding sequence (CDS), were annotated as putative pseudogenes. The graphical representation of *O. quimilo* circular annotated plastome was created in OGDRAW (Lohse et al., 2013). To visualize changes in gene order and content, we compared the *O. quimilo* assembly with the canonical gene order of *P. oleracea* plastome via multiple whole genome alignment using MAUVE (default options, assuming colinearity; Darling et al., 2004). Boundaries between the IRa IRb, LSC, SSC and putative inversions were visually checked in Geneious using an *in silico* approach adapted from Oliver et al. 2010.

### 2.3. Comparative chloroplast genome sequences analyses

The newly annotated plastome of *Opuntia quimilo*, with one of the inverted repeats (IRa) manually stripped to avoid data duplication, was then used for a reference guided assembly on the trimmed reads from all other taxa using Geneious mapper with a medium-low sensitivity iterating up to 5 times (adapted from Ripma et al 2014). Each of the assembles mapped had a majority threshold consensus sequence generated and an annotation transferred from the *O. quimilo* reference, and manually adjusted. To identify highly variable regions across the subfamily, the 17 assembled Opuntioideae chloroplast genome sequences were compared using mVista (Frazer et al., 2004) in Shuffle-LAGAN alignment mode (Brudno et al. 2008) using the annotated plastome of *O. quimilo* as a reference. We also used the full chloroplast genome sequence alignment (see below) to calculate nucleotide diversity values (π) to detect highly variable sites among Opuntioideae chloroplast genome sequences. DNA polymorphisms analysis was performed on DnaSP v.6.10 (Rozas et al. 2017) using the sliding window analysis with step size of 200 bp and window length of 800 bp. Assembly maps of raw read coverages from Geneious mapper of each taxon to the *O. quimilo* plastome were also used to visualize and compare the gene content of the chloroplast genome sequences across the subfamily.

### 2.4. Phylogenetic analyses and informative regions

The assembled chloroplast genome sequences were aligned using MAFFT v. 7 with an automatic strategy search for algorithm selection (Katoh & Standley 2013), using 200PAM scoring matrix and an open gap penalty of 1.53 (offset value 0.123). The alignment was manually examined for misaligned areas following a similarity criterion (Simmons, 2004). Sequence portions that contained gaps and/or ambiguities across more than 80% of the taxa were stripped using the “Mask Alignments” feature in Geneious. Phylogenetic inference was performed using Maximum Likelihood implemented in RAxML 8.2.4 (Stamatakis, 2014) in the CIPRES Portal (Miller et al. 2010) with GTR+G model employed for the entire sequence. Support values were estimated implementing 1,000 bootstrap replicates.

In order to identify and rank highly phylogenetically informative regions in the Opuntioideae plastomes, we split the full plastome alignment in protein coding sequences (cpCDS – pseudogenes were included here), non-coding sequences (cpNCDS) and intergenic spacers (cpIGS) using the annotated *O. quimilo* plastome. Each individual marker (cpCDS, cpNCDS, cpIGS) was extracted from the above-mentioned alignment, and a Maximum Likelihood tree was inferred with RAxML (GTR+G model, 100 bootstrap replicates). For each marker, we report the number of variable sites, number of parsimony informative sites (PIS), mean sequence distance (under K80 model), alignment length, mean sequence length, mean bootstrap support and distance to the full chloroplast genome sequence tree (RF distance; Robinson & Foulds, 1981). The metrics were retrieved using functions of the R packages *ape* and *phangorn* (Paradis, Claude & Strimmer, 2004; Schliep, 2011). Markers were ranked by phylogenetic information using a weighted mean of relative values of the following metrics: number of variable sites (weight = 1), mean bootstrap (weight = 2) and distance to the full plastid tree (weight = 3). We designed primer pairs for the top 5 markers identified in the previous step with suitable size for PCR amplification (∼800–900 bp). Primers flanking the target regions were designed with Primer3, using the default settings (Rozen & Skaletsky, 2000). All metrics reported, as well as primer design, were considered only for the ingroup (the 17 Opuntioideae chloroplast genome sequences).

## 3. Results

### 3.1 DNA sequencing

Runs on Illumina HiSeq X resulted in 227,003,814 reads from 20 samples (17 Opuntioideae and 3 outgroups), between 5,624,110 and 20,219,350 reads per sample, for a mean read number of 11,350,190 sequences. Reads per sample following quality control were between 5,360,990 and 19,863,298 with a mean post-quality control read pool number of 11,084,834. The GC content of the raw reads ranged from 37.4% to 40.6% with a mean of 38.45% and following quality control were between 36.9% and 40% with a mean of 38%. Detailed results with the number of raw reads, post-quality control and %GC content per taxa are presented in Table S1.

### 3.2. Opuntia quimilo plastome

The complete chloroplast genome of *Opuntia quimilo* was sequenced, assembled, annotated and deposited in GenBank with the accession number MN114084. The length of the *Opuntia quimilo* plastome is 150,374 bp, including a 101,475 bp LSC region, a 4,115 bp SSC region and a 22,392 bp of two IR (IRa and IRb) region (Figure 1, Table 1). A total of 701,318 reads were assembled, with an average organelle depth of 844x. The GC content varies from 33% in the SSC, to 35.5% in LSC and 39.6% in the IR regions (Table 1).

**Figure 1.**
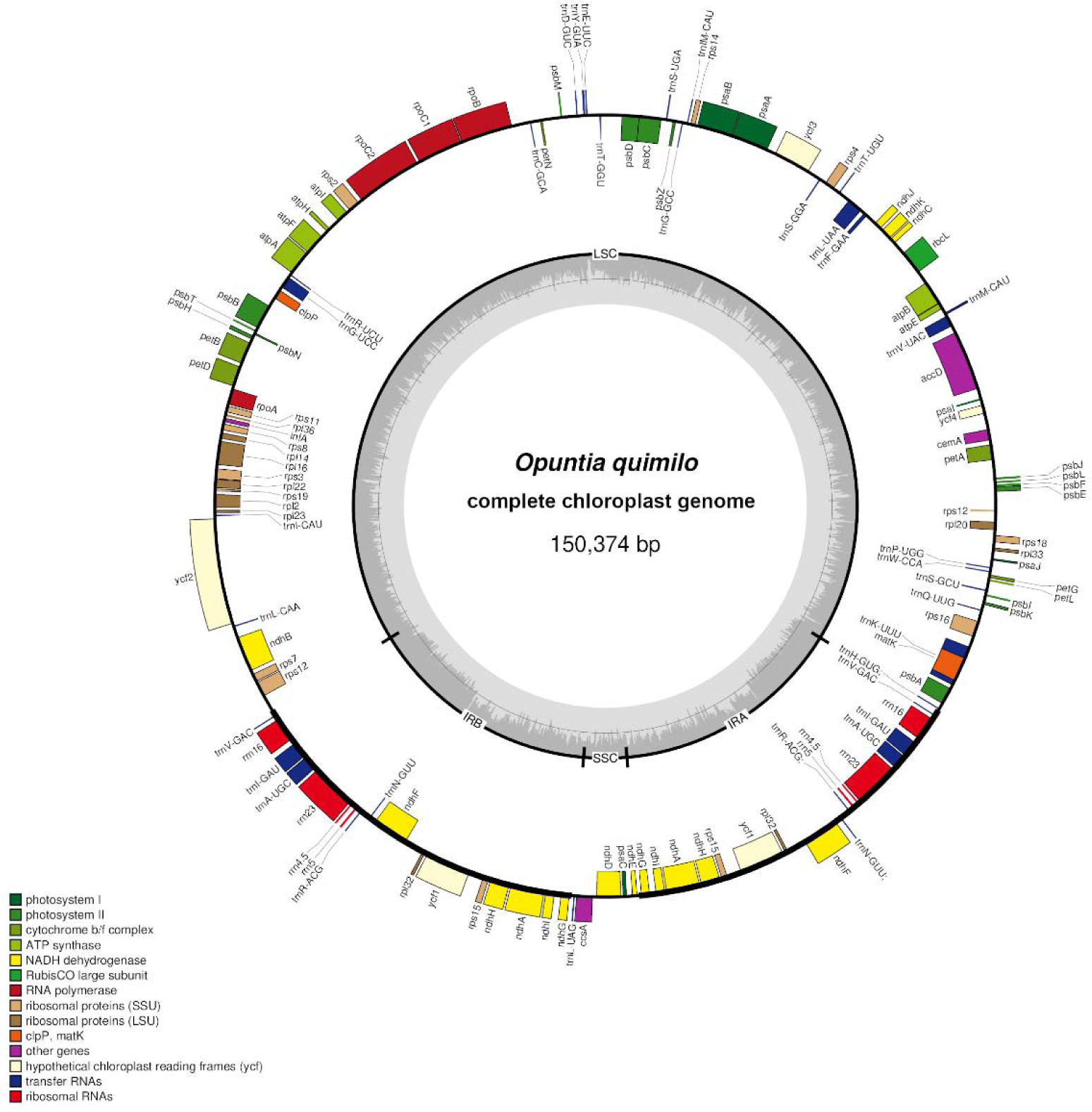
Circular map of chloroplast genome of Opuntia quimilo with annotated genes. The genes transcribed clockwise are shown inside of the circle whereas genes transcribed counter clockwise are shown outside of the circle. The borders of chloroplast genome are defined with LSC, SSC, IRa and IRb. The dashed gray color of inner circle shows the GC content.

**Table 1.** Chloroplast genome composition of *Opuntia quimilo*.

| **Region** | **Size (bp)** | **GC (%)** | **Genes** | **CDS** | **tRNA** | **rRNA** |
| --- | --- | --- | --- | --- | --- | --- |
| **Genome** | 150.374 | 36.6 | 130 (5) | 87 (5) | 35 | 8 |
| **LSC** | 101.475 | 35.5 | 91 (4) | 67 (4) | 24 | 0 |
| **SSC** | 4.115 | 33 | 5 | 4 | 1 | 0 |
| **IRa** | 22.392 | 39.6 | 17 (1) | 8 (1) | 5 | 4 |
| **IRb** | 22.392 | 39.6 | 17 (1) | 8 (1) | 5 | 4 |

The number in parentheses indicates pseudogenes ( $\Psi$ ) identified.

The *de novo* assembly of Geneious assembler produced 1,000 contigs; of these, 988 were higher than 1,000 bp in length from a minimum length of 1,026 bp to a maximum of 283,150 bp. MegaBLAST search founds one consensus plastid contig of 128,909 bp that includes the full chloroplast sequence with two putative inverted repeats assembled as a single IR unit (∼22 kb). The GetOrganele and NOVOPlasty pipelines both yielded one plastid contig of 150,374 bp with the same gene content, order and structure as the plastid contig of the Geneious assembler, except for the two inverted repeats that were interleaved by the LSC and SSC on the first ones while in the Geneious contig was merged as one IR.

The *Opuntia quimilo* plastome encodes 87 protein-coding genes (CDS), 35 transfer RNA genes (tRNA) and 8 ribosomal RNA (rRNA) genes, totaling 130 genes (Table 1 and 2). Five canonical CDS from angiosperm chloroplast genomes were annotated as putative pseudogenes (Ψ) based on their structure: *accD*, *rpl16*, *rps16*, *ycf1* and *ycf2*. Four of them are located in the LSC, and one (*ycf1*) in the IRs. Duplicated CDS in the IRs includes *ndhA, ndhF*, *ndhG*, *ndhH*, *ndhI*, *rpl32*, *ycf1*(Ψ) and *rps15*; and all four rRNA genes and five of the 35 tRNA are duplicated in the IR regions. The *O. quimilo* plastome includes 16 intron-containing genes, of which 12 contain one intron (*atpF*, *ndhA, ndhB, rps12, rps16, rpl16, rpoC1, petB, petD, trnA^UGC^*, *trnI^GAU^*, *trnK^UUU^*, *trnV^UAC^*, *trnL^UAA^*, *trnG^UCC^*), while one gene contains two introns (*ycf3*); the *clpP* gene has lost its two introns, reduced to an exon of 609 bp.

**Table 2.**
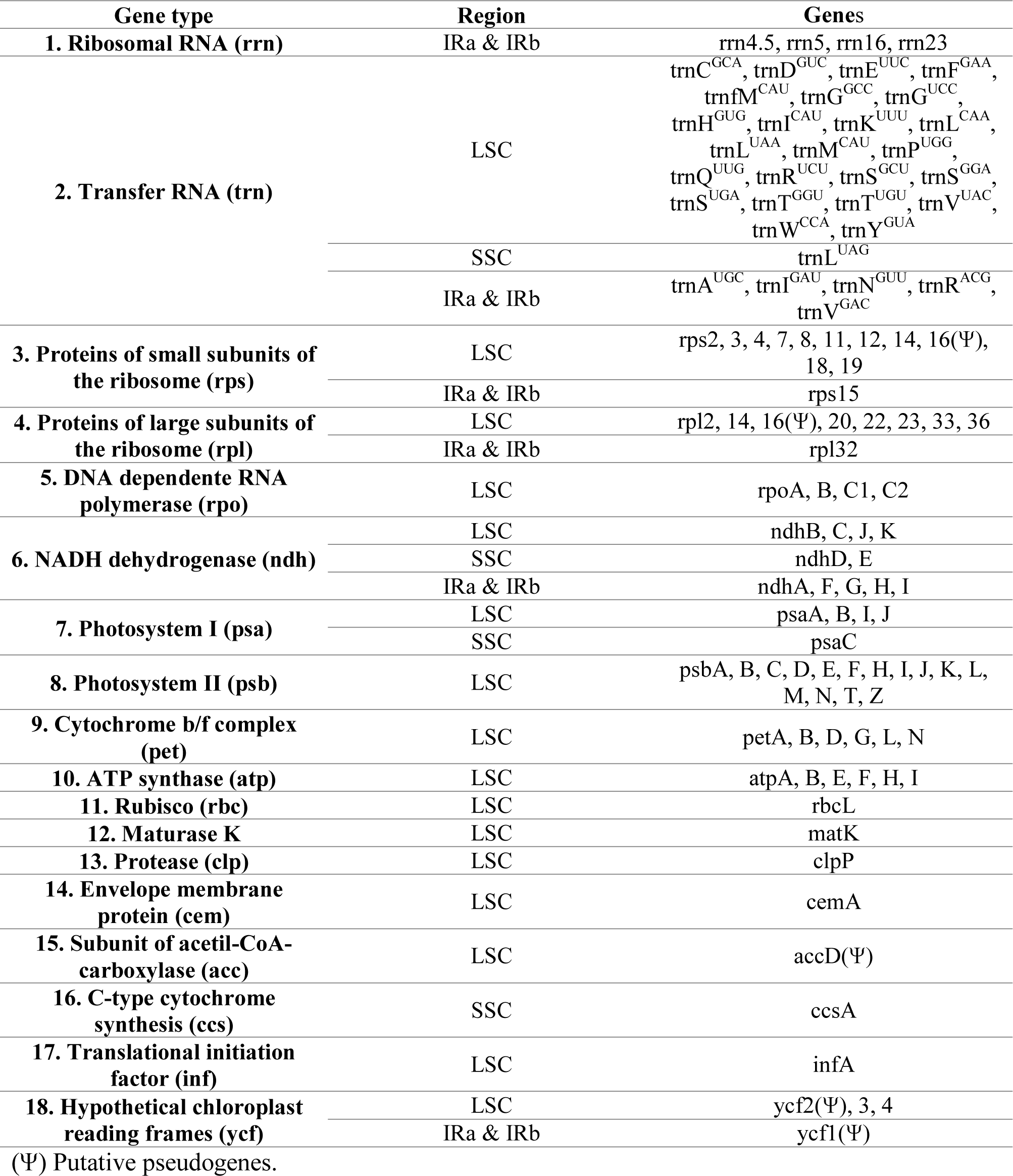
Structural and functional gene composition of *Opuntia quimilo* chloroplast genome

The LSC of the *Opuntia quimilo* plastome appears to have experienced an expansion, with surprisingly 101 kb, while the SSC was shown to have exceptional reduction (4 kb). The LSC contains 24 tRNA genes and 67 CDS, and the SSC contains an unique tRNA gene (*trnL^UAG^*), and four CDS: *ccsA*, *ndhE*, *ndhD* and *psaC* (Figure 1 and 2; Table 1 and 2). A total of eight genes (*ndhB*, *rpl2, rpl23, rps7, rps19, trnI^CAU^*, *trnL^CAA^* and *ycf2*) that are usually reported occurring in the IR regions of canonical angiosperm plastomes are apparently present as unique genes - not repeated - in the LSC region of the *O. quimilo* plastome (Fig. 2, region V). On the contrary, seven genes (*ndhA, ndhF, ndhG, ndhH, ndhI, rpl32* and *rps15*), usually from the SSC, are duplicated into the IR regions of *O. quimilo* plastome (Fig. 2, orange genes).

**Figure 2.**
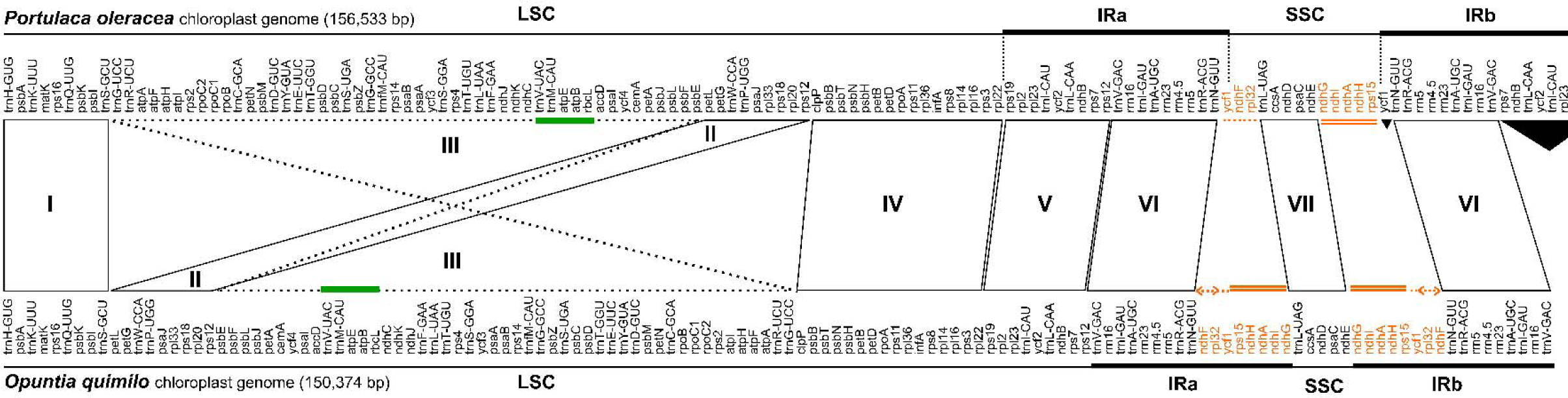
Plastid genome structure and gene order in Opuntia quimilo compared with purslane (Portulaca oleracea). Purslane has the canonical order typical of most angiosperms. For simplicity, the circular map has been linearized. Green line highlights the trnMCAU-rbcL synapomorphic inversion of Cactaceae, which in O. quimilo also includes the trnVUAC gene. Regions I, IV, V, VI and VII are colinear in both plastomes. Region II is colinear but is translocated in the O. quimilo plastome, while region III is inverted and translocated. Region V comprise the genes that are typically in the IR region but are translocated to the large single copy in O. quimilo. Genes highlighted in orange are those typically found in the SSC but transferred to the IR region in O. quimilo. Orange dashed line indicate the double inversion on the ycf1-rpl32 genes, placing ycf1 gene adjacent to rpl32. Black triangles represent duplicated genes present in purslane but absent in O. quimilo; LSC = large single-copy region; SSC = small single-copy region; IR = Inverted repeat.

When compared to the canonical angiosperm chloroplast genome of *Portulaca oleracea*, two block translocations in the LSC are present in the *O. quimilo* plastome: the first (Fig. 2, region II) is a simple colinear translocation of nine genes (Fig. 2, region II); while the second one is a big block inversion and translocation comprising 50 genes within the *trnG^UCC^-psbE* region (Fig. 2, region III). Inside that block (region III), the putative synapomorphic inversion of cacti encompassing the *trnM-rbcL* genes is confirmed for Cactaceae, but in the *O. quimilo* plastome this inversion also encompassed the *trnV^UAC^* gene (Fig. 2, green bars). Further gene order is mainly colinear (Fig 2., regions I, IV, V, VI, VII), except for the rearrangement comprising the SSC genes that were transferred to the IRs regions, including a double inversion on the *ycf1*-*rpl32* region, placing *ycf1* gene adjacent to *rpl32* (Fig. 2, orange genes).

### 3.3. Reference-guide assemblies and comparative chloroplast sequences analyses

The reference-guided assembles of the remaining Opuntioideae and outgroup taxa to the *Opuntia quimilo* plastome (one inverted repeat stripped) mapped an average of 616,615 reads with a mean genome depth of 721x (Table S3). The consensus sequence length varied between 126,925 bp (*Pereskiopsis diguettii*) to 129,181 bp (*Tacinga palmadora*) and the GC content between 35.8% (*Pterocactus gonjianii*) to 36.3% (*Austrocylindropuntia cylindrica* and *Cylindropuntia bigelovii*) (Table S3).

**Table 3.**
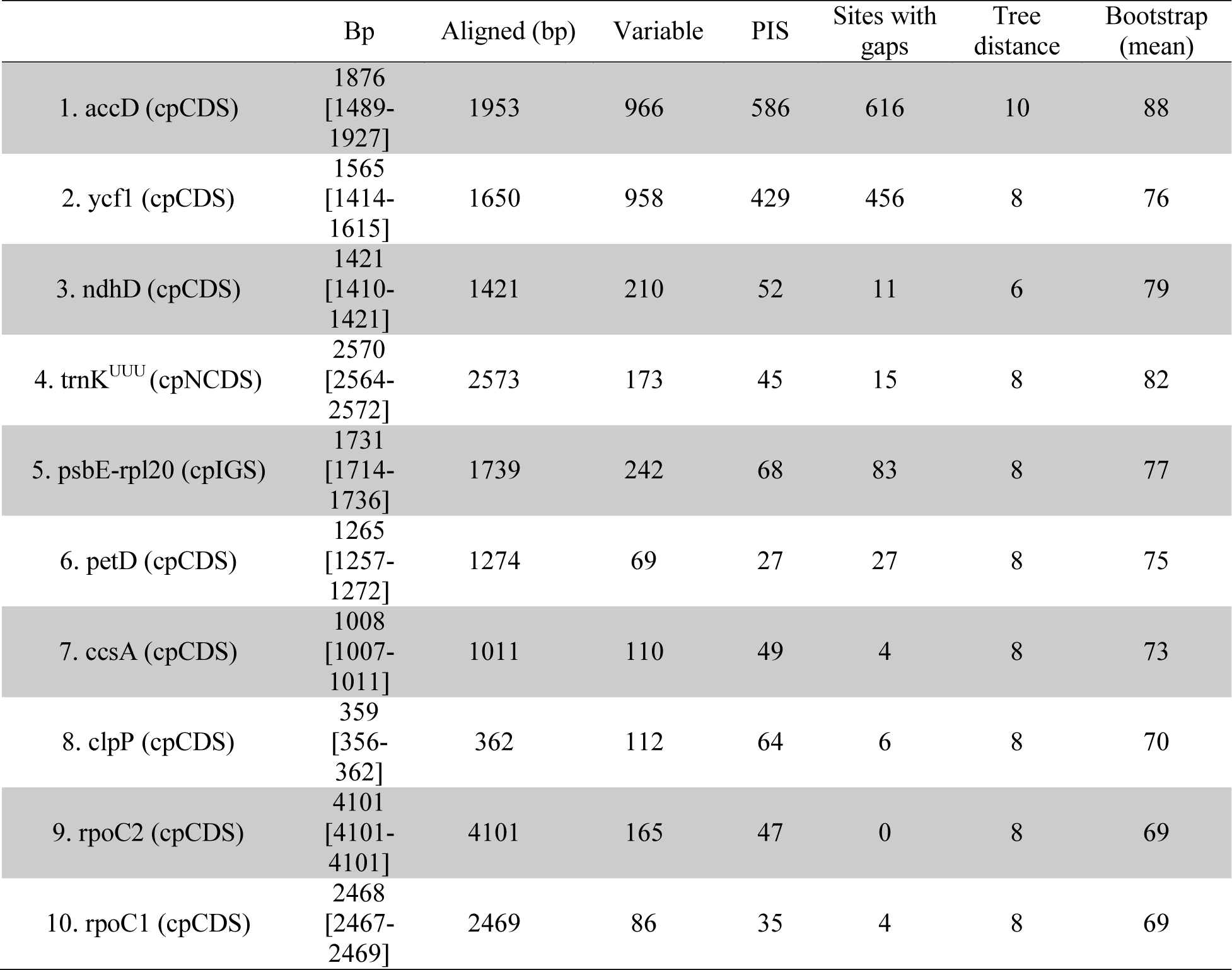
Summary statistics for the top 10 markers. Markers are ranked by phylogenetic information based on a weighed mean of relative values of number of variable sites (weight = 1), mean bootstrap (weight = 2) and distance to the full plastid tree (weight = 3).

|  | Bp | Aligned (bp) | Variable | PIS | Sites with gaps | Tree distance | Bootstrap (mean) |
| --- | --- | --- | --- | --- | --- | --- | --- |
| 1. accD (cpCDS) | 1876<br>[1489-1927] | 1953 | 966 | 586 | 616 | 10 | 88 |
| 2. ycf1 (cpCDS) | 1565<br>[1414-1615] | 1650 | 958 | 429 | 456 | 8 | 76 |
| 3. ndhD (cpCDS) | 1421<br>[1410-1421] | 1421 | 210 | 52 | 11 | 6 | 79 |
| 4. trnK<sup>UUU</sup> (cpNCDS) | 2570<br>[2564-2572] | 2573 | 173 | 45 | 15 | 8 | 82 |
| 5. psbE-rpl20 (cpIGS) | 1731<br>[1714-1736] | 1739 | 242 | 68 | 83 | 8 | 77 |
| 6. petD (cpCDS) | 1265<br>[1257-1272] | 1274 | 69 | 27 | 27 | 8 | 75 |
| 7. ccsA (cpCDS) | 1008<br>[1007-1011] | 1011 | 110 | 49 | 4 | 8 | 73 |
| 8. clpP (cpCDS) | 359<br>[356-362] | 362 | 112 | 64 | 6 | 8 | 70 |
| 9. rpoC2 (cpCDS) | 4101<br>[4101-4101] | 4101 | 165 | 47 | 0 | 8 | 69 |
| 10. rpoC1 (cpCDS) | 2468<br>[2467-2469] | 2469 | 86 | 35 | 4 | 8 | 69 |

Pairwise comparison of divergent regions within the Opuntioideae chloroplast genome sequences using mVISTA with *O. quimilo* as a reference revealed both striking conserved and divergent regions across the chloroplast genomes sequences (Figure 3). Overall, the alignment uncovered sequence divergence across assemblies, suggesting that chloroplast genome sequences are not conserved. Divergences were observed both in noncoding regions and coding regions. Among coding regions (cds), non-conserved regions were frequent on genes of the *ndh* gene suite (i.e. *ndhA, ndhD, ndhE*, *ndhF*, *ndhG*, *ndhH, ndhI*, *ndhJ*) as well *clpP*, *ycf3* and particularly highlighted on *ycf1*, *ycf2* and *accD* genes (Fig. 3). Ten noncoding regions show substantial divergence, being all intergenic spacers: *ndhE-psaC, rpl32-ndhF, trnV^GAC^-rps12, psbB-clpP, rpoB-trnC^GCA^, psbM-trnD^GUC^, trnT^GGU^-psbD, psbE-rpl20, ndhC-rbcL* (Fig. 3).

**Figure 3.**
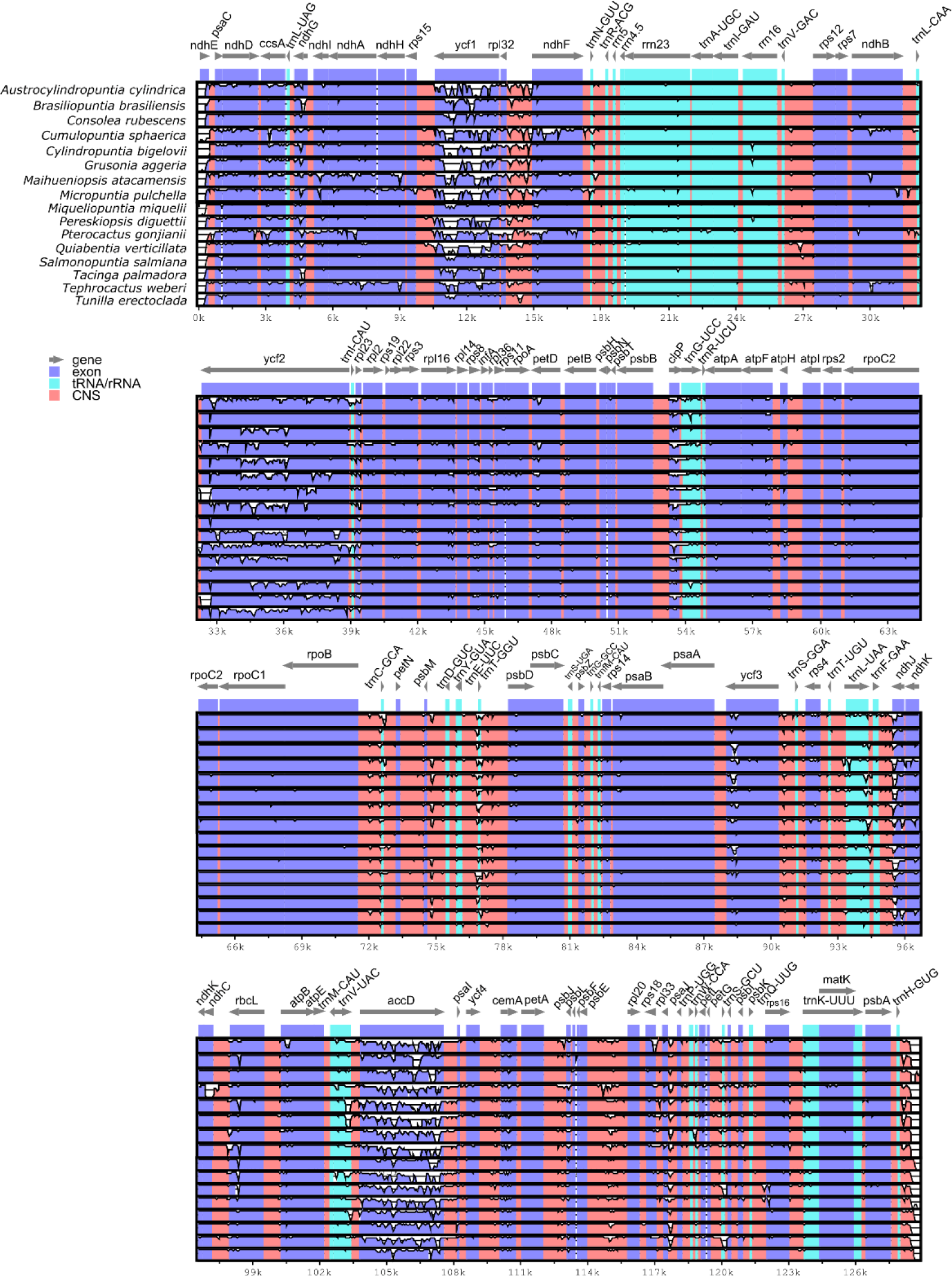
Visualized alignment of the Opuntioideae chloroplast genome sequences (one IR stripped) with annotations using mVISTA. Each horizontal lane shows the graph for the sequence pairwise identity with Opuntia quimilo as reference. The x-axis represents the base sequence of the alignment and the y-axis represents the pairwise percent identity within 50– 100%. Grey arrows represent the genes and their orientations. Dark-blue boxes represent exon regions; light-blue boxes represent tRNA and rRNA regions; red boxes represent non-coding sequence (CNS) regions.

The nucleotide diversity values (π) within the 17 Opuntioideae chloroplast genome sequences ranged from 0. 00191 to 0.18551, with a mean value of 0.02201, indicating the sequences as highly variable. Three major regions were identified as hypervariable (π > 0.1), which comprises *ycf1* and *accD* genes and an intergenic spacer *rpl32-ndhF* (Figure 4); while six regions were observed as moderately-variable (π > 0.05), those being four genes: *ycf2*, *ccsA*, *clpP* and *trnL^UAA^*; and two intergenic spacers *rps18-rpl33* and *trnF^GAA^-ndhJ* (Figure 4).

**Figure 4.**
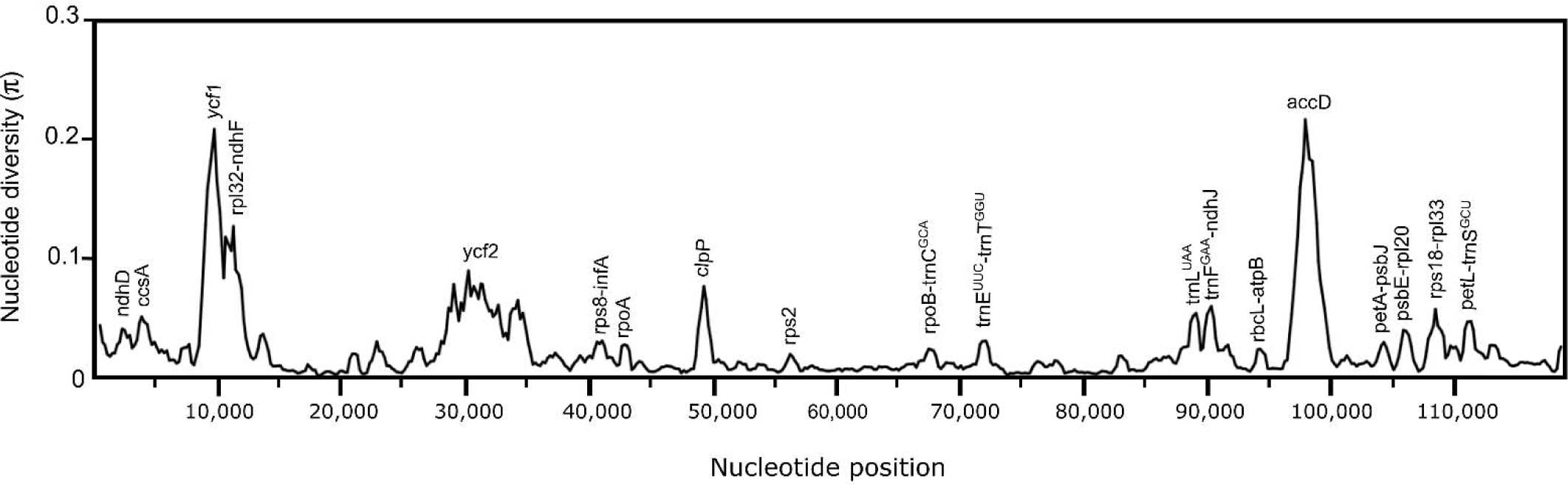
Nucleotide diversity graphs of the 17 Opuntioideae chloroplast genome sequences from the sliding windows analysis performed in DnaSP (windows length: 800 bp, step size: 200 bp). The x-axis re alignment the base sequence of the alignment, and the y-axis represents the nucleotide diversity (π value). Each variation hotspot for the chloroplast genome sequences of Opuntioideae alignment is annotated on the graph.

Reference-guided assembled maps of Opuntioideae and outgroups to the *Opuntia quimilo* chloroplast genome as a reference revealed regions with extremely low coverage or even gaps across different taxa (Figure 5). The regions highlighted with this feature are related with genes of the *ndh* suite, *ycf1*, *ycf2* and *accD*, suggesting gene loss, transfer to nuclear genomes and/or pseudogenization (Figure 5). Several members of Opuntioideae appear to have missing *ndh* genes in their chloroplast genome (*Micropuntia, Maihueniopsis, Pterocactus, Tephrocactus*), especially in the Tephrocacteae clade. However, no clear pattern of low coverage among these regions was recovered within Opuntioideae clades, indicating putative homoplasious events.

**Figure 5.**
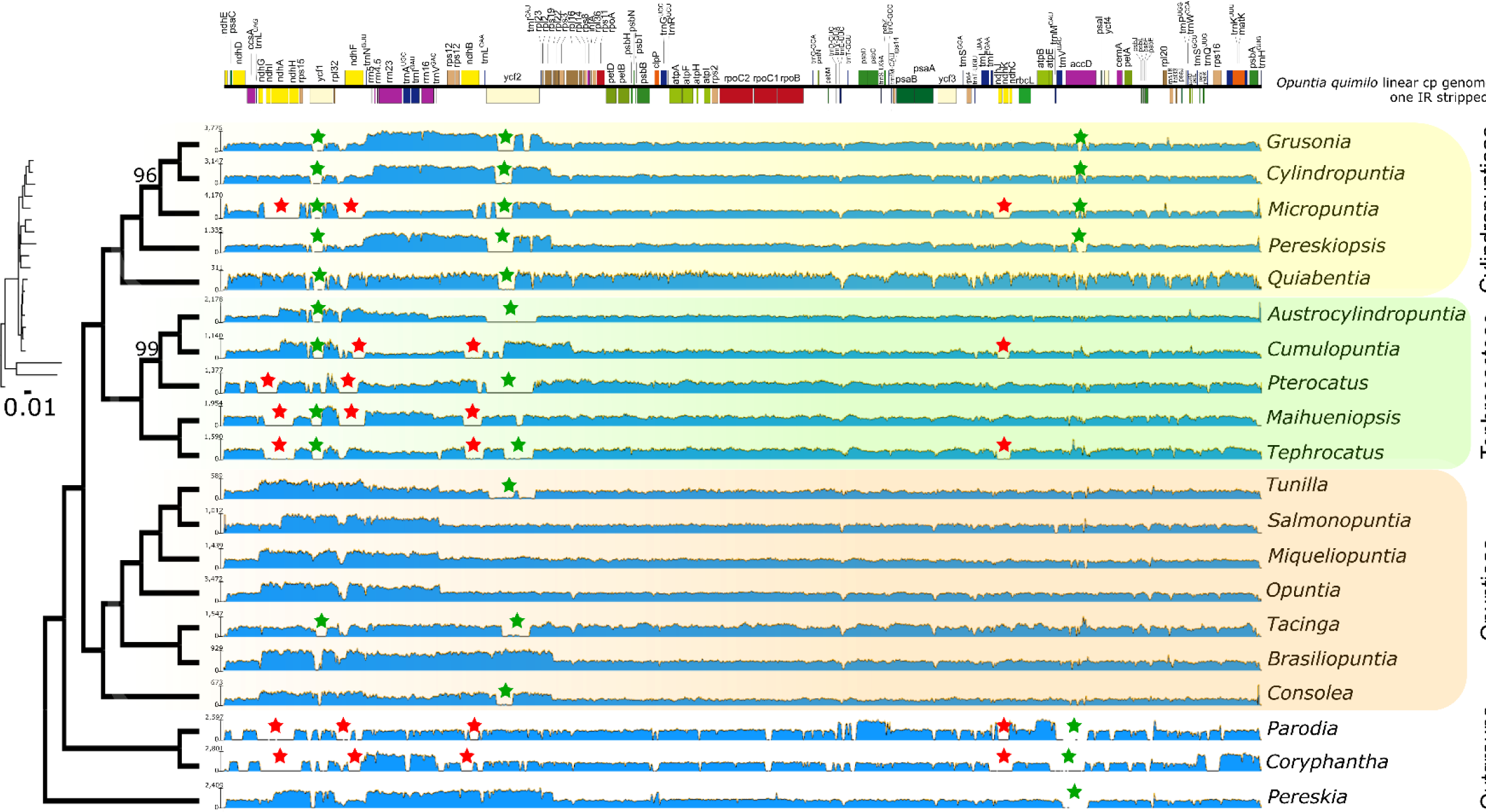
Maximum likelihood phylogenetic tree from RAxML analysis transformed in cladogram with the phylogram represented in small size with substitution rate scaled. All nodes have total bootstrap values (bs = 100) with exception for those that are shown above the branch. Each tip is represented with the assembly map of raw read coverages from Geneious mapper to the Opuntia quimilo chloroplast genome (one IR stripped, represented on the top with annotated genes). Red stars represent low coverage mapping and putative losses associated with the ndh gene suite; green stars represent partial low coverages associated with putative pseudogenization of ycf1, ycf2 and accD genes. Tribe Opuntieae is highlighted in orange, Tephrocacteae in green and Cylindropuntieae in yellow.

### 3.4. Phylogenetic analyses and informative regions

The full chloroplast genome sequences resulted in an alignment of 118,930 bp with 86,484 identical sites (72.7%), a pairwise identity of 94.5% and 8,694 distinct alignment patterns. There are 8,922 parsimony informative sites (PIS) and 11,509 sites with gaps. Maximum Likelihood analyses resolved a well-supported Opuntioideae (bs = 100), with three major subclades currently circumscribed as tribes, i.e., Opuntieae, Cylindropuntieae and Tephrocateae (Fig. 5). Opuntieae, consisting of the seven genera *Consolea*, *Brasiliopuntia*, *Tacinga*, *Opuntia, Miqueliopuntia*, *Salmonopuntia* and *Tunilla*, was resolved as sister to Tephrocacteae (*Thephrocatus*, *Maihueniopsis*, *Pterocactus*, *Cumulopuntia* and *Austrocylindropuntia*) and Cylindropuntieae (*Quiabentia*, *Pereskiopsis*, *Micropuntia*, *Grusonia* and *Cylindropuntia*) clade. All nodes had high bootstrap support values (bs= 100), except by two nodes, which were still higher than 90 (Fig. 5).

The summary statistics for all markers (cpCDS, cpNCDS, cpIGS) are presented in Table S4. A list of the top 10 markers ranked by phylogenetic information considering topological distance to the plastome tree, mean bootstrap support and number of parsimony informative sites is given in Table 3. All single marker phylogenies presented some disagreement to the plastome tree (RF tree distance ranging from 6 to 28). Bootstrap support ranged from 0 to 89 (mean = 37), and number of PIS from 0 to 619 (mean = 25). Primer pair sequences for PCR amplification are provided for the top 5 markers with suitable Sanger sequencing size (max ∼900 bp) in Table 4.

**Table 4.**
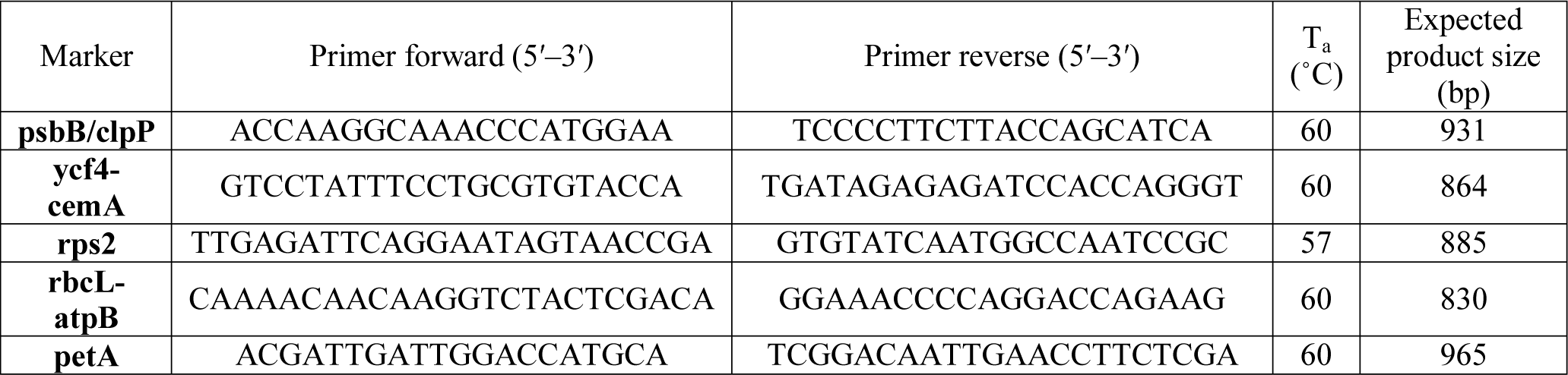
Primer pair sequences for the identified top five highly informative markers across the 17 chloroplast genome sequences of Opuntioideae.

## 4. Discussion

### 4.1 Insights from chloroplast genome assemblies in Opuntioideae and Cactaceae

The first chloroplast genome of a species of *Opuntia* is here reported. Although the bulk of its gene content is not far from canonical angiosperms plastomes, it deviates in some cases from the typical chloroplast genome structure, showing: (i) an expansion of the LSC incorporating genes that are typically in the IRs; (ii) a reduction of the SSC translocating some common genes of the SSC into the IR region; and (iii) at least one massive translocation with an inversion of a block of genes in the LSC (Figure 2). Part of the content of the IRs in the *O. quimilo* plastome remained remarkably constant, including all four rRNA and five rRNA genes that are nearly universally reported in IRs of land plants (Mower & Vickrey 2018).

Successive expansion–contraction events or even multiple contractions have been recurrently reported as one of the main ways of developing structural changes across Angiosperm plastomes (Downie & Jansen 2015; Daniell et al 2016; Fonseca & Lohmann 2017; Mower & Vickrey 2018) and may also be one way in which genes are translocated to different regions of the genome, as suggested in adzuki bean (Perry et al. 2002). The atypical reduction of the SSC (∼ 4 kb), reported here for the *O. quimilo* plastome, has also been noticed in *Viviana marifolia* (Francoaceae, Geraniales), and a slightly similar reduced size for SSC (∼6 kb) have been inferred for the ancestral chloroplast genome of Geraniaceae (Weng et al. 2013). A partial deletion of SSC region has also been reported in two hemiparasitic *Taxillus* (Loranthaceae) species resulting in a ∼6 kb region with only two genes (Li et al. 2017), and the smallest SSC hitherto reported is for the hemiparasitic *Pedicularis ishidoyana* (Orobanchaceae), with only 27 bp (Cho et al. 2018). A model to explain the major rearrangements observed in the *Lamprocapnos spectabilis* (Papaveraceae) plastome, involving at least six IR boundary shifts and five inversions resulting in a SSC of just 1,645 bp with a partial ndhF gene, was recently provided by Part et al. (2018). The SSC contains most *ndh* genes, and previous studies have shown that boundary shifts of the IR and SSC regions are correlated with transformations of *ndhF* and *ycf1* genes (Kim et al. 2015; Logacheva et al. 2014; Li et al. 2017).

The *Opuntia quimilo* plastome reinforces some different putative structural synapomorphies that have been reported in Caryophyllales. For example, the loss of the *rpl2* intron, previously suggested to be absent throughout the Centrospermae (Palmer et al. 1988), is supported in our study and others newly assembled plastomes in Caryophyllales (Yao et al. 2019). The *trnM-rbcL* inversion is again recovered in the *O. quimilo* plastome, although also involving the *trnV^UAC^* gene, as in *Cylindropuntia bigelovii* (Majure et al. 2019), providing further support for this inversion as a synapomorphy in the family. Additionally, Sanderson et al. (2015) and Solorzano et al. (2019), inspecting plastomes of Cactoideae, reported a gene orientation of *ycf2*-*trnL^CAA^*-*ycf1* in the SSC as a synapomorphy of Cactoideae. Our results corroborate this observation, since this feature is not present in the *O. quimilo* plastome, strengthening this gene order as a putative synapomorphy for Cactoideae. On the other hand, the *ycf1-rpl32-ndhF* orientation, reported in the *Cylindropuntia bigelovii* chloroplast sequence (Majure et al. 2019), is recovered in the *O. quimilo* plastome and is here suggested as a putative synapomorphy for Opuntioideae.

Besides our *de novo* assembly and chloroplast genome conception of *Opuntia quimilo*, our reference-guide assemblies and comparative analyses revealed insights for plastome rearrangements across disparate Opuntioideae. The differences of depth and coverage among specific chloroplast genes may suggest that gene structure or presence can vary over species in Opuntioideae, as have been observed in other Cactaceae, specifically Cactoideae (Sanderson et al. 2015; Solarzano et al. 2019). The putative independent losses of several *ndh* genes in all Cactoideae plastomes assembled hitherto, such as the saguaro cactus and several *Mammillaria* species, can be also inferred for our Cactoideae outgroups sampled (*Parodia magnifica* and *Coryphantha macromeris;* Fig. 5, red stars). Likewise, some members of Cylindropuntieae and Tephrocacteae (*Micropuntia*, *Cumulopuntia*, *Pterocactus*, *Maihueniopsis* and *Tephrocactus*) also likely experienced independent losses of several genes of the *ndh* suite in their chloroplast genomes, although this was not so for tribe Opuntieae, where those genes were found to be intact (Fig. 5, red stars).

The major plastid regions marked by pseudogenization in the *Opuntia quimilo* plastome (*ycf1*, *ycf2* and *accD*) are visually highlighted as non-conserved regions both in reference-guide maps (Fig. 5, green stars), as in the mVista alignment across Opuntioideae (Fig. 3). These regions are also emphasized as with hyper or moderate variability regarding the nucleotide diversity values (Fig. 4). All genes here reported as pseudogenes in the *O. quimilo* plastome (*accD*, *rpl16*, *rps16*, *ycf1* and *ycf2*) have also been reported as pseudogenes in the *Mammillaria* plastomes (Solórzano et al. 2019), while the *accD* was described as a pseudogene in *Carnegia gigantea* (Sanderson et al. 2015). Pseudogenization of these genes has been repeatedly reported across different angiosperm lineages, such as Malpighiales, Campanulales, Ericales, Poales, Solanales, Geraniales, Santalales and Myrtales (Harris et al. 2013; Haberle et al. 2008; Fajardo et al. 2013; Weng et al. 2013; Bedoya et al. 2019; Li et al. 2017; Cui et al. 2019; Machado et al. 2017). Even though these genes have been identified with essential functions beyond photosynthesis and retained in the plastome of most embryophytes (Drescher et al., 2000; Kuroda and Maliga, 2003; Kode et al., 2005; Kikuchi et al., 2013; Parker et al., 2014; Dong et al., 2015), there are several other plants where these genes are missing from the chloroplast genome (Kim, 2004; Magee et al., 2010; Lei et al., 2016; Graham et al., 2017). The pseudogenization or loss of the *accD*, *rpl22* and several genes of the *ndh suite* from the plastids has been reported to be a consequence of them being transferred to the nuclear genome (Jansen et al., 2011; Jansen and Ruhlman 2012; Sanderson et al. 2015; Cauz-Santos et al., 2017; Liu et al. 2016). Plastid gene transfer to the nucleus remains to be examined in *O. quimilo* and related Opuntioideae.

Several regions highlighted as hyper or moderately variable regarding the nucleotide diversity values across Opuntioideae chloroplast sequences (i.e., *accD*, *ycf1*, *clpP, petD, rpl32* and *ccsA*) have been reported to be putatively under positive selection in some lineages, such as Brassicaceae, Bignoniaceae, Rutaceae, Orchidaceae, Geraniaceae and Poaceae (Hu et al. 2015; Weng et al. 2016; Dong et al. 2018; Carbonell-Caballero et al. 2015; Ruhlman and Jansen 2018; Thode & Lohmann 2019; Park et al. 2017; Piot et al. 2018). Positive selection may come into play in response to environmental changes (Piot et al. 2018). For example, the *accD* gene, which encodes the β -carboxyl transferase subunit of acetyl-CoA carboxylase, is an essential and required component for plant leaf development (Kode et al. 200), and it is suggested to have played a pivotal role in the adaptive evolution of orchids (Dong et al. 2018). The signatures of positive selection under *accD* gene observed in Brassicaceae and Campanulaceae have also indicated that this gene may have been repeatedly involved in the adaption to specific ecological niches during the radiation of eudicotyledonous plants (Rousseau-Gueutin et al. 2013; Hu et al. 2015). Considering the harsh environment that cacti display their fitness already expressed in its peculiar morphology and physiology, further studies should be carried to investigate the putative relation of chloroplast rearrangement – such as pseudogenization, loss of genes, translocation and inversion – with ecological features.

### 4.2. Phylogenetic relationship of Opuntioideae tribes

The plastome phylogeny of Opuntioideae strongly resolves three major clades that are currently circumscribed as tribes Opuntieae, Tephrocacteae and Cylindropuntieae (Fig. 5 and Fig. S1). Three previously described tribes (Austrocylindropuntieae, Pterocacteae and Pereskiopsideae) (Doweld 1999; Wallace and Dickie, 2002) are nested within these more broadly circumscribed tribes and thus have no real logical or practical taxonomic use (Hunt 2011). Our results partially recovered previous topologies (Griffith & Porter 2009; Ritz et al. 2012; Walker et al. 2018) and agreed with those of Majure et al. (2019) but differed from those by the strong support for the relationships among clades, as well the expanded sampling including all accepted genera in Opuntioideae. As our study is still based on one sample per genus, future studies including more taxa per genus should be carried out across the subfamily to further test the relationships here recovered.

Tribe Opuntieae is sister to the rest of Opuntioideae, and the most diverse and widespread clade among Opuntioideae, consisting of seven accepted genera (Majure & Puente 2014, Guerrero et al. 2018). *Consolea* Lem., an endemic tree-like cactus of the Caribbean Islands and neighboring areas, is sister to the rest of Opuntieae, which consists of two subclades: (i) one comprising *Brasiliopuntia* (K. Schum.) A. Berger + *Tacinga* Britton & Rose; and the other comprising (ii) *Opuntia* (L.) Mill. + (*Miqueliopuntia* Frič ex F. Ritter + (*Salmonopuntia* P.V. Heath + *Tunilla* D.R. Hunt & Iliff.)). Morphological evolution in Opuntieae are discussed in Majure and Puente (2014), although our results presented a slightly different topology for the relationships of *Consolea* and (*Miqueliopuntia* (*Salmonopuntia* + *Tunilla*)).

Cylindropuntieae and Tephrocacteae are sister tribes, each one comprised of five genera (Fig. 5). Cylindropuntieae are primarily represented by genera that occur in the western North American desert regions (*Cylindropuntia* (Engelm.) F.M. Knuth, *Grusonia* F. Rchb. & K. Schum. and *Micropuntia* Daston), which formed a well-supported subclade, but they also contain two genera that are found in tropical dry forest of Mexico/Northen Central America (*Pereskiopsis* Britton & Rose) and South America (*Quiabentia* Britton & Rose). Tribe Pereskiopsideae (Doweld 1999), previous described to only accommodate the leafy *Pereskiopsis*, is nested within Cylindropuntieae and is redundant, and thus unnecessary. Deeper relationships within Cylindropuntieae were recently untangled using a phylogenomic approach and dense sampling, revealing biogeographic patterns as well as characters evolution (Majure et al. 2019).

Tephrocacteae is a South American clade adapted to diverse climatic conditions over a wide area of the southern Andes and adjacent lowlands (Ritz et al. 2012; Guerrero et al. 2018; Las Peñas et al. 2019). The tribe includes morphologically diverse species from geophytes and cushion-plants to dwarf shrubs, shrubs or small trees (Anderson 2001); and probably geophytes and cushion-forming species evolved several times from shrubby-like precursors (Ritz et al. 2012). Tribes Austrocylindropuntieae and Pterocacteae (Wallace & Dick 2002), described to circumscribe *Autrocylindropuntia* + *Cumulopuntia* and *Pterocactus*, respectively, are both nested in the Tephrocacteae, as amplified by Hunt (2011), and their use is mostly redundant.

### 4.3. Phylogenetically informative regions

Our plastome survey for phylogenetically informative markers revealed a list of potentially highly informative plastid markers for Sanger-based phylogenetic studies in Opuntioideae (Table S4). The top 10 markers in our cpCDS dataser are: *accD, ycf1*, *ndhD, petD, ccsA, clpP, rpoC1, rpoC2*, including just one intron (the *trnK* intron comprising the *matK* gene – *trnK/matK*) and one intergenic spacer (*psbE-rpl20*) (Table 3). However, two of the better ranked markers (*accD* and *ycf1*) are putative pseudogenes and must be treated apart from traditional protein coding genes.

From the top 10 markers ranked in our list, just one (*trnK/matK*) has been used in more than one phylogenetic study in cacti (Nyffeler 2002; Edwards et al. 2005; Korotova et al. 2010; Demaio et al. 2011; Arakaki et al. 2011; Bárcenas et al. 2011; Hernández-Hernández et al. 2011; Hernández-Hernández et al. 2014; Ritz et al. 2012; Bárcenas 2016; Luna-Vargas et al. 2018); while Majure et al. (2012) has used partial sequences of the *ycf1* gene. The others top 10 markers have been previously used under phylogenomic approaches in cacti (Arakaki et al. 2011; Majure et al. 2019).

Although the majority of the top 10 markers here reported have not been used in phylogenetic studies of cacti, the relationship of several other groups has been inferred with some of these markers. For example, the *accD* gene, combined with other plastid regions including *rpoC1*, was employed for phylogenetic inference of *Crocus* (Iridaceae), *Coptis* (Ranunculaceae) and Orchidaceae genera (Peternsen et al. 2008; Guo et al. 2012; He et al. 2014). However, *accD* intergenic spacers, such as *rbcL-accD* and *accD-psaI,* have been much more widely used across disparate groups (Barfuss et al. 2005; Miikeda et al. 2006; Reginato et al. 2010; Sun et al. 2012; Michelangeli et al. 2012). The *ycf1* gene appears to be moderately used (Gernandt et al. 2009; Guo et al. 2012; Majure et al. 2012; Whitten et al. 2013; Shi et al. 2013; Dastpak et al. 2018), and increasingly reported to be a useful marker in phylogenetics inferences (Neubig et al. 2009; Neubig and Abbott 2010; Dong et al. 2012; Thomson et al. 2018), and the most promising plastid DNA barcode of land plants (Dong et al. 2015). The *petD* intron has been used (Löhne et al. 2007; Borsh et al. 2009; Worberg et al. 2007; Scataglini et al. 2013), but in our analysis the entire gene was used (exon + intron) showing phylogenetic utility. The *ccsA* gene seems to be underexplored as a phylogenetic marker (Marx et al. 2010; Peterson et al. 2012) but was already suggested as convenient for phylogenetic inferences (Logacheva et al. 2007). The *rpoC1* and *rpoC2* genes have been occasionally used together as markers (Liston and Wheeler 1994; Kulshreshtha et al. 2004) or combined with other markers (GPWS 2001; Zhang et al. 2011; Downie et al. 2000; Guo et al. 2012) yielding satisfactory results. The *rpoC2* gene was recently found as the best performing marker to recover with high levels of concordance the “accepted tree” of the angiosperm phylogeny (Walker et al. 2019). The *ndhD* gene seems to be scarcely used for phylogenetic inference (Panero and Funk 2002), while the intergenic spacer of *psbE-rpl20* genes has never been used individually to our knowledge.

Eight of the top 10 markers are more than 900 bp, indicating that longer genes are superior for phylogeny reconstruction, as previous suggested by Walker et al. (2019), although they may require internal primer designing for complete Sanger’s sequencing. A list of the top 10 markers with less than 900 bp is reported (Table S5), and primer pair design for the top five is provided in Table 4. The *clpP* gene (∼ 359 bp) is the best ranked under this criterium (< 900 bp), and the intergenic spacer *psbB-clpP* (∼ 547 bp) is also ranked in the top 10 list of this list, thus, the pair primer design included them as one marker (*psbB/clpP*). Many of the top 10 markers listed with less than 900 bp have been occasionally used in phylogenetic studies across disparate plant lineages with variable resolution, but never in cacti. The *clpP* gene is usually employed with its introns (Stefanović et al. 2009; Lam et al. 2016), and the *psbB-clpP* intergenic spacer has been increasingly reported as useful marker (Loera et al. 2012; Särkinen and George 2013; Prince 2015). The intergenic spacer *ycf4-cemA* has been used for phylogenetic studies in Asteraceae and Poaceae genera (Zhang et al. 2016; Ekenas et al. 2007) and the *rps2* gene in Orobanchaceae and Ephedraceae (McNeal et al. 2013; Manen et al. 2004; Loera et al. 2012). The *petA* gene appears to be rarely used (Tsumura et al. 1996), while the intergenic spacer *petA-psbJ* has been employed across various groups (Jaramillo et al 2008; Guo and Wilson 2013; Wang et al. 2014), including Opuntioideae (Majure et al. 2012). The intergenic spacer *ycf3-trnS* has been used in phylogenetic inferences in Asteraceae and Euphorbiaceae (Vilatersana et al. 2010; Montes Moreno et al. 2010; Barres et al. 2011), and the *rps3* gene in Poaceae, Verbenaceae and families of Asparagales (Peterson et al. 2010; Marx et al. 2010; Kim et al. 2012). The intergenic spacer *ndhE-psaC* appears to have not yet been used as a marker for phylogenetic studies, but it was also reported as useful for phylogenetic and phylogeographic studies in Liliaceae (Lu et al. 2017). The intergenic spacer *ndhC-rbcL* is putatively exclusive of Cactaceae, resulting from the *trnV-rbcL* inversion, and its phylogenetic utility must be further investigated, at least in clades where *ndh* genes are present.

Chloroplast markers have been used for testing evolutionary relationships among plants for the past 30 years (Gitzendanner et al. 2018). While the assumption that these markers are evolving as a single unit without recombination, routine analyses have concatenated data producing highly supported phylogenies that have been underlying the current classification of angiosperms (APG, 2016). However, as here reported, no marker as a single unit (gene tree) recovered the same topology of the plastome inference (concatenated tree), and even within the top 10 markers listed, some showed high values of discordance (Table 3 and Table S5). Such results discourage and call attention to phylogenetic approaches based on one or few markers. Recent studies have explored gene tree conflicts in plastome-inferred phylogenies and incongruence between gene trees and species trees in plastid genes (Walker et al. 2019; Gonçalves et al. 2019). Gonçalves et al (2019) emphasized the importance of considering variation in phylogenetic signal across plastid genes and the exploration of plastome data to increase accuracy of estimating relationships; they also revealed that phylogenies inferred with multispecies coalescent (MSC) methods are accurate with plastome matrices and should be considered in future phylogenomic investigations. Walker et al. (2019) highlighted that most genes are largely uninformative and are unlikely to misguide plant systematics. However, the concatenating of plastid genes without some level of scrutiny can mislead branch length estimation (Walker et al. 2019). The causes of discordant topologies across gene trees from chloroplast genome still needs to be better investigated. The main explanations include systematic errors (e.g. poor modeling, stochastic events) or more biologically meaningful processes, such as heteroplasmic recombination that have been invoked to explain discordance in disparate plant clades (Huang et al. 2001; Marshall et al. 2001; Bouillé et al. 2011; Walker et al. 2019).

## 5. Conclusions

Chloroplast genomes have long been considered conserved among land plants, but recent generation of thousands of plastomes through NGS has illuminated that this is not always the case. Cactaceae are no exception to variations that have been observed. Previous plastomes of cacti have shown to have lost one copy of the inverted repeat regions and several genes of the *ndh* gene suite, as well as to possess divergent inverted repeat regions and the smallest chloroplast genome known for an obligately photosynthetic angiosperm. We showed that the *Opuntia quimilo* plastome also presents deviations of canonical angiosperm plastomes with an expansion of the LSC incorporating genes that are typically in the IRs, a reduction of the SSC translocating some common genes of the SSC into the IR region, and one massive translocation with an inversion of a block of genes in the LSC. Our reference-guided assemblies across Opuntioideae allowed us to infer putative independent losses of some *ndh* genes across disparate taxa of the subfamily. We did not find synapomorphic plastome features within Opuntioideae clades, thus, we hypothesize that putative rearrangements across the subfamily are from homoplasious events. Further analyses should be carried out to address how ecological drivers and morphological traits of cacti may be related with positive selection of genes and the common rearrangements in chloroplast genomes that have been reported in the family. Phylogenetic analyses of chloroplast genome sequences strongly support Opuntioideae and its three tribes: Opuntieae, Cylindropuntieae and Tephrocacteae. Since computational and budget limitations are still a bottleneck to deal with high throughput data, especially in developing countries, a list of highly informative plastid markers is presented for future use, and several top ranked markers have not been used in phylogenetic studies of cacti. Nonetheless, gene trees discordances between plastome markers must be carefully considering while inferring phylogenies in this remarkable group of plants.

## Acknowledgments

The authors thanks to Grant Godden (FLMNH) for provide valuable comments about chloroplast assemblers; Urs Eggli (Sukkulenten-Sammlung Zürich) for provide important bibliography; and to Ricardo Reis and Josimar Külkamp (Jardim Botânico do Rio de Janeiro, Brazil) for helping providing samples. MK is grateful to the American Society of Plant Taxonomists (ASPT), Cactus and Succulent Society of America (CSSA), International Association for Plant Taxonomy (IAPT) and IDEA WILD for supporting part of the research here reported; the Brazilian National Council for Scientific and Technological Development (Conselho Nacional de Desenvolvimento Científico e Tecnológico – CNPq, process number 141523/2017-4) for his PhD scholarship, and the PDSE/CAPES for support his period as Visiting Researcher at the Florida Museum of Natural History (FLMNH, UF, USA, Process 88881.186882/2018-01). TTSC acknowledge CNPq (grant number: 304506/2016-8).

## Funding

Partial funding for this project was provided by the Desert Botanical Garden (Arizona, USA), start-up funds from the University of Florida to L.C. Majure, the National Science Foundation (DEB 1735604), and the ‘Coordenação de Aperfeiçoamento de Pessoal de Nível Superior – Brasil’ (CAPES) - Finance Code 001.

## Author Contributions

**MK**: Conceptualization, Methodology, Validation, Formal analysis, Investigation, Data Curation, Writing - Original Draft, Writing - Review & Editing, Visualization, Project administration, Funding acquisition; **MR**: Methodology, Software, Validation, Formal analysis, Writing - Review & Editing; **TTSC**: Writing - Review & Editing; **LCM**: Methodology, Validation, Investigation, Resources, Data Curation, Writing - Review & Editing, Supervision, Funding acquisition.

---

**Table S1.** Taxa sampled in this study, raw and post–quality control read numbers, GC content and voucher information. Vouchers contain collector name, number and herbarium repository in parentheses or the accession number from the Desert Botanical Garden Living collection (https://dbg.org/research-conservation/living-collections/).

| **Taxon** | **Raw Read N°** | **Read N° Post-QC** | **%GC** | **Voucher** |
| --- | --- | --- | --- | --- |
| *Austrocylindropuntia cylindrica* | 7724354 | 7362548 | 37.4 | *Baker19306* (ASU) |
| *Brasiliopuntia brasiliensis* | 5624110 | 5360990 | 39.6 | *Köhler 412* (ICN) |
| *Consolea rubescens* | 6227612 | 6038548 | 37.4 | *Majure 3320* (FLAS) |
| *Coryphantha macromeris* | 8294236 | 7997166 | 40.6 | *Majure 7128* (FLAS) |
| *Cumulopuntia sphaerica* | 8245847 | 8106104 | 37.6 | DBG1993-0190 |
| *Cylindropuntia bigelovii* | 16876574 | 16431880 | 39.2 | *Baker 18286* (ASU) |
| *Grusonia aggeria* | 16876574 | 16084111 | 39.5 | *Majure 5651* (DES) |
| *Maihueniopsis atacamensis* | 17489908 | 17322547 | 37.4 | DBG1993-0194 |
| *Micropuntia pulchella* | 20219350 | 19863298 | 39 | *Majure 6095* (DES) |
| *Miqueliopuntia miquelii* | 15182404 | 15065674 | 37.6 | DBG1993-0205 |
| *Opuntia quimilo* | 16137088 | 15814820 | 38.5 | DBG2003-0111 |
| *Parodia magnifica* | 5792409 | 5786100 | 38.3 | DBG1993-0275 |
| *Pereskia sacharosa* | 8781612 | 8526372 | 38.4 | DBG2014-2153 |
| *Pereskopsis diguettii* | 8789752 | 8650470 | 38 | DBG2018-0128 |
| *Pterocactus gonjianii* | 9941712 | 9809046 | 38.9 | DBG1997-0260 |
| *Quiabentia verticillata* | 9223832 | 8893090 | 38.1 | *Kimnach 2803* (HNT) |
| *Salmonopuntia salmiana* | 6667776 | 6425092 | 38.8 | *Köhler 515* (ICN) |
| *Tacinga palmadora* | 18511428 | 18182592 | 39.2 | DBG1997-0392 |
| *Tephrocactus weberi* | 13461630 | 13300339 | 37.8 | DBG2012-0428 |
| *Tunilla erectoclada* | 6935606 | 6675910 | 37.8 | DBG2018-0144 |

**Table S2.** Reference-guide assemblies of Opuntioideae taxa to the *Opuntia quimilo* plastome.

| Taxon | Chloroplast |  |  |  |  |
| --- | --- | --- | --- | --- | --- |
|  | Reads Mapped | Genome Coverage | Consensus Sequence Length | % GC | GenBank Submission |
| *Austrocylindropuntia cylindrica* | 551,356 | 642 | 127,734 | 36.3 |  |
| *Brasiliopuntia brasiliensis* | 386,272 | 449 | 128,560 | 36.1 |  |
| *Consolea rubescens* | 191,291 | 221 | 128,464 | 36.2 |  |
| *Coryphantha macromeris* | 817,912 | 982 | 128,341 | 36.2 |  |
| *Cumulopuntia sphaerica* | 333,795 | 393 | 127,844 | 36 |  |
| *Cylindropuntia bigelovii* | 1,159,529 | 1355 | 128,307 | 36.3 |  |
| *Grusonia aggeria* | 1,307,477 | 1534 | 127,410 | 36.2 |  |
| *Maihueiniopsis atacamensis* | 661,251 | 775 | 128,476 | 36.2 |  |
| *Micropuntia pulchella* | 1,189,650 | 1404 | 127,876 | 36.1 |  |
| *Miqueliopuntia miquelii* | 544,818 | 638 | 128,543 | 36.1 |  |
| *Parodia magnifica* | 810,368 | 954 | 127,406 | 36.2 |  |
| *Pereskia sacharosa* | 919,347 | 1075 | 128,319 | 35.9 |  |
| *Pereskiopsis diguettii* | 449,688 | 523 | 126,925 | 36 |  |
| *Pterocactus gonjiani* | 481,070 | 552 | 127,885 | 35.8 |  |
| *Quiabentia verticillata* | 170,524 | 197 | 128,319 | 36.3 |  |
| *Salmonopuntia salmiana* | 392,127 | 453 | 128,810 | 36.1 |  |
| *Tacinga palmadora* | 598,318 | 699 | 129,181 | 36.1 |  |
| *Tephrocactus weberi* | 548,088 | 632 | 128,302 | 36.1 |  |
| *Tunilla erectoclada* | 202,807 | 236 | 128,352 | 36.1 |  |

**Table S3.**
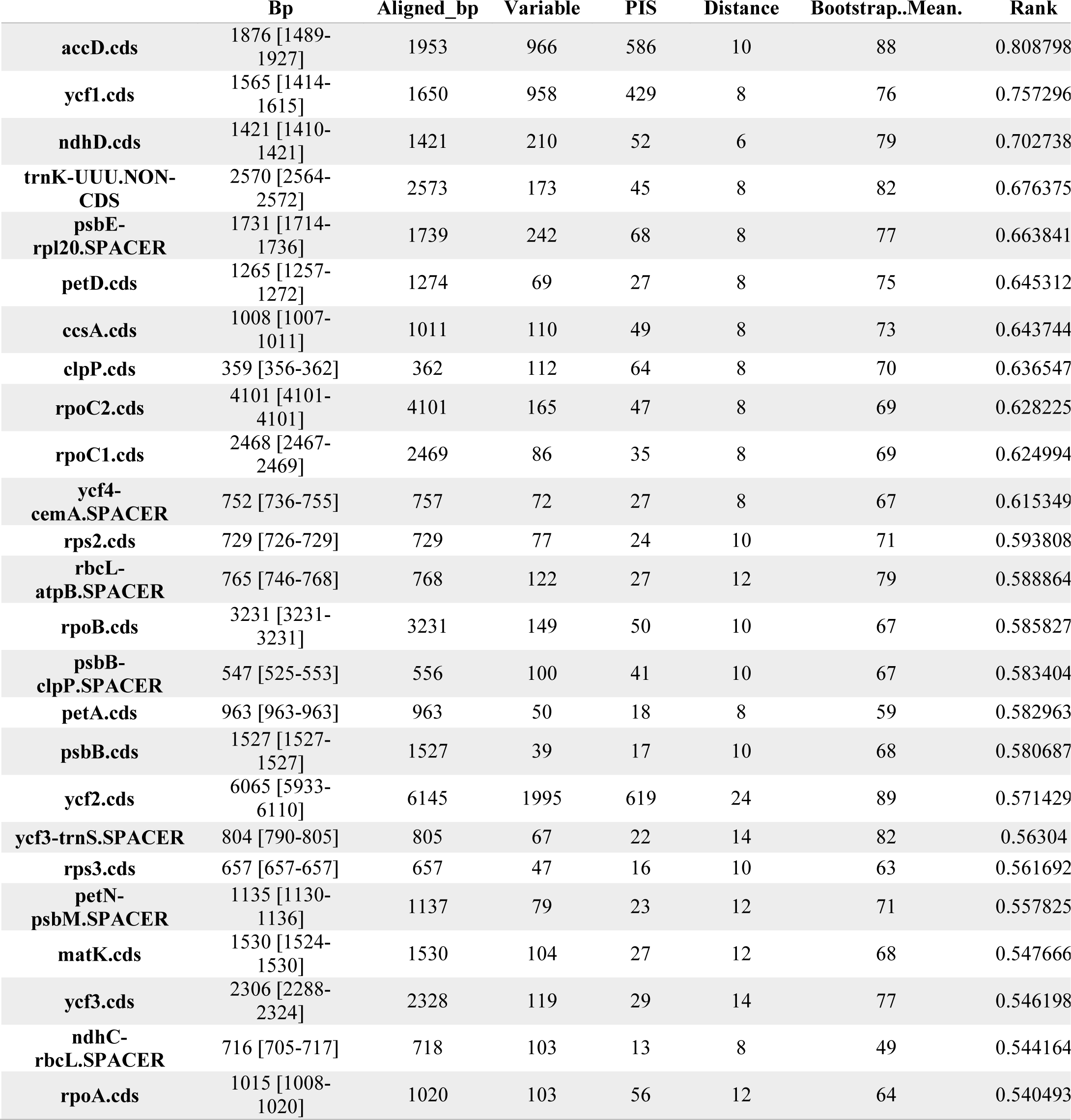
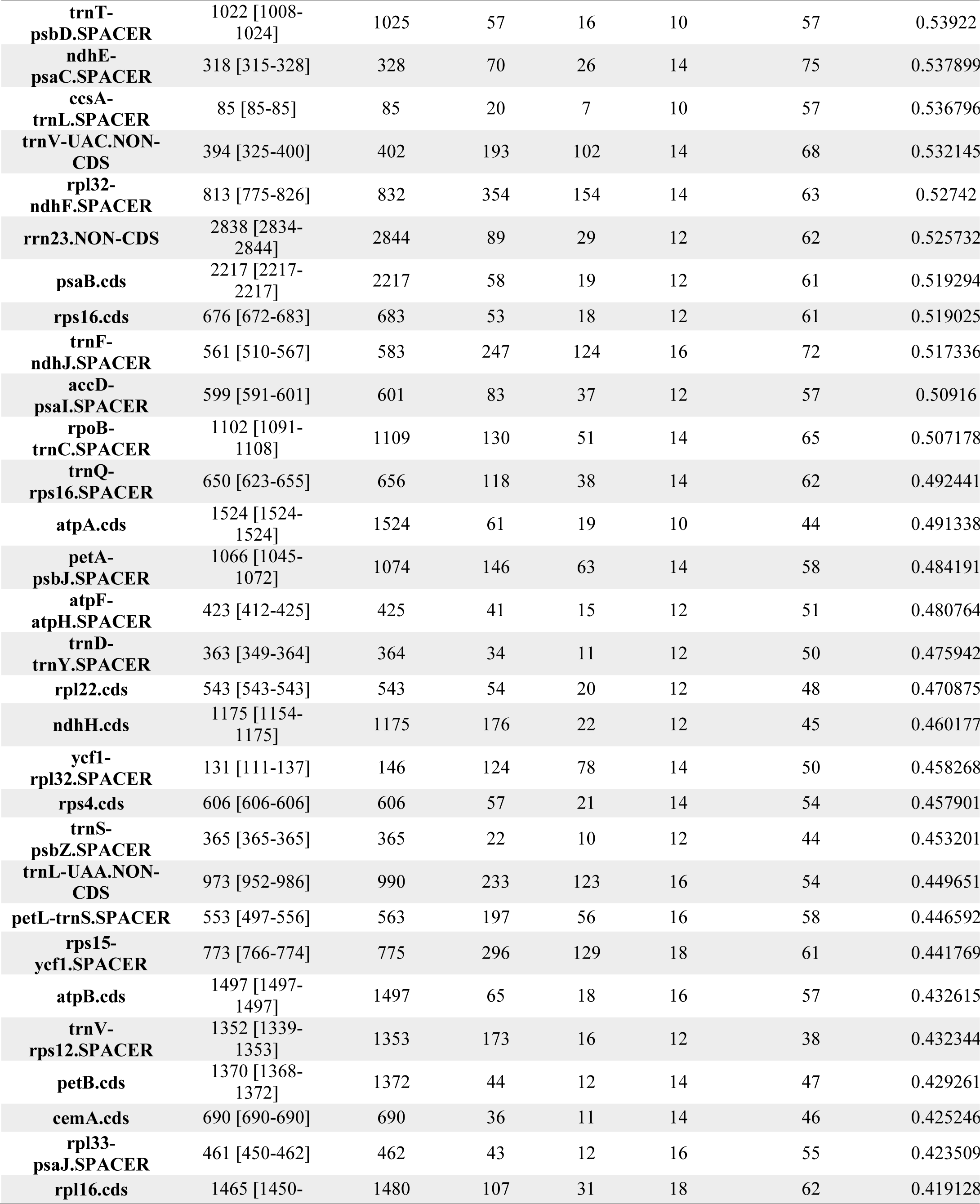
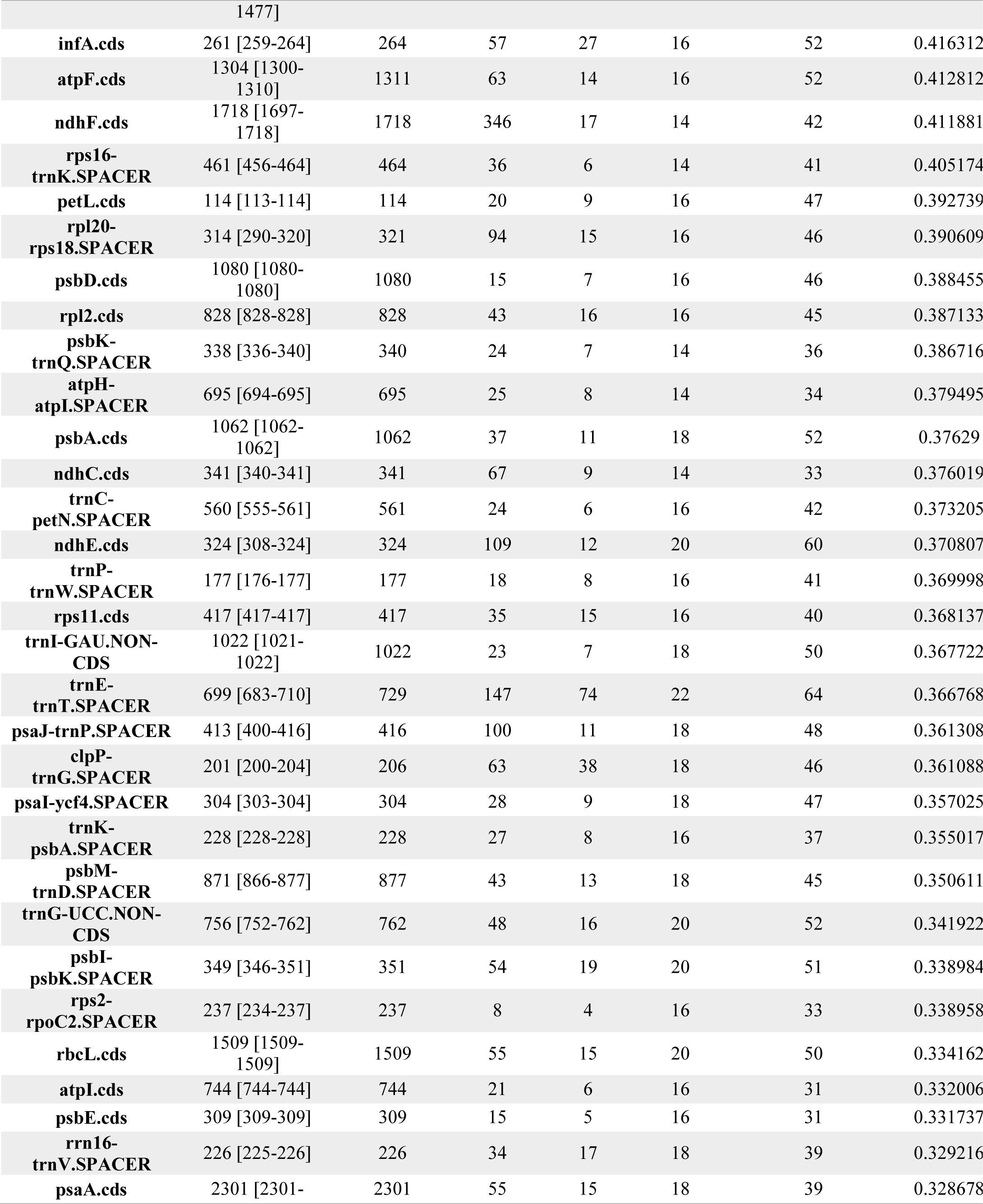
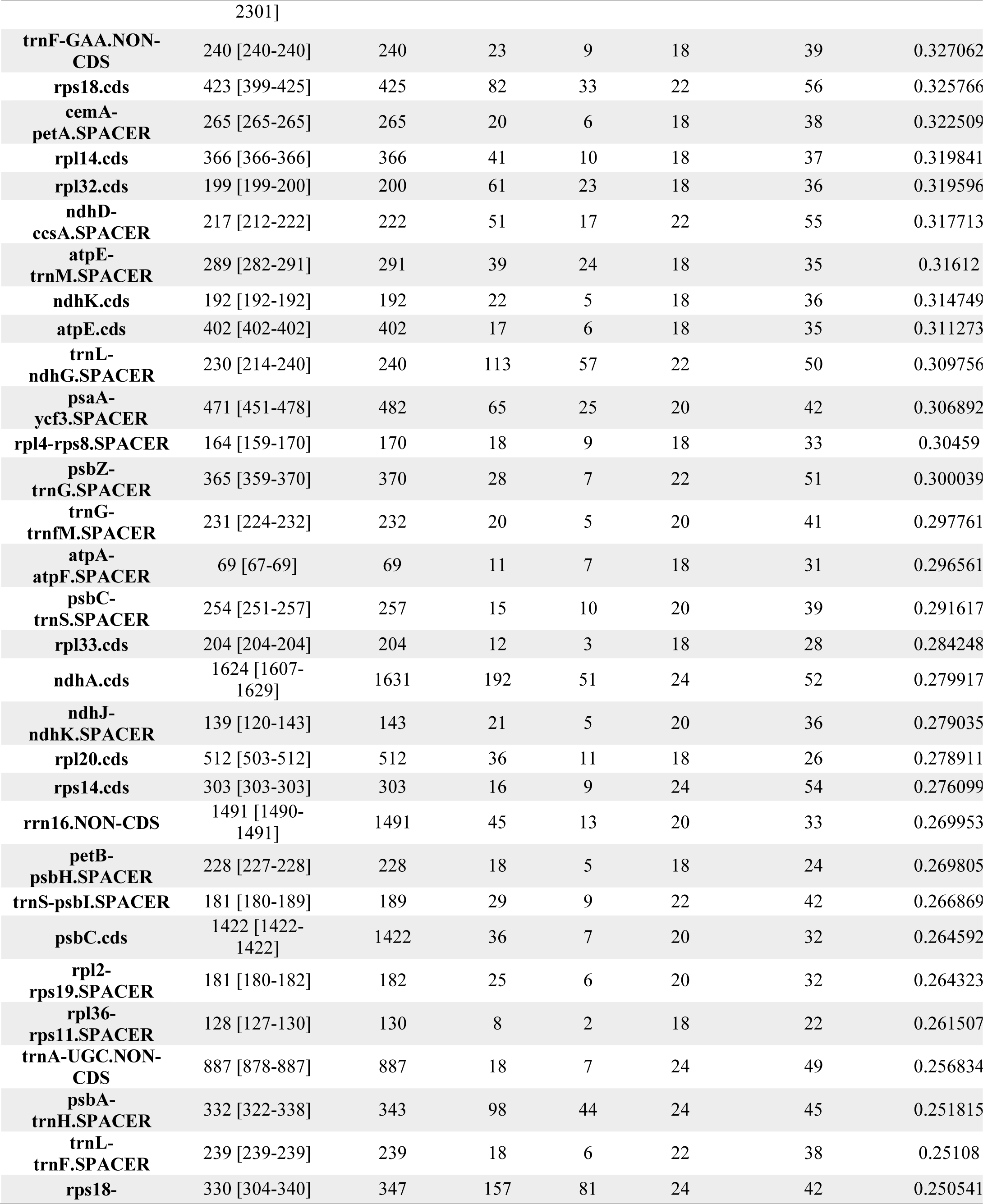
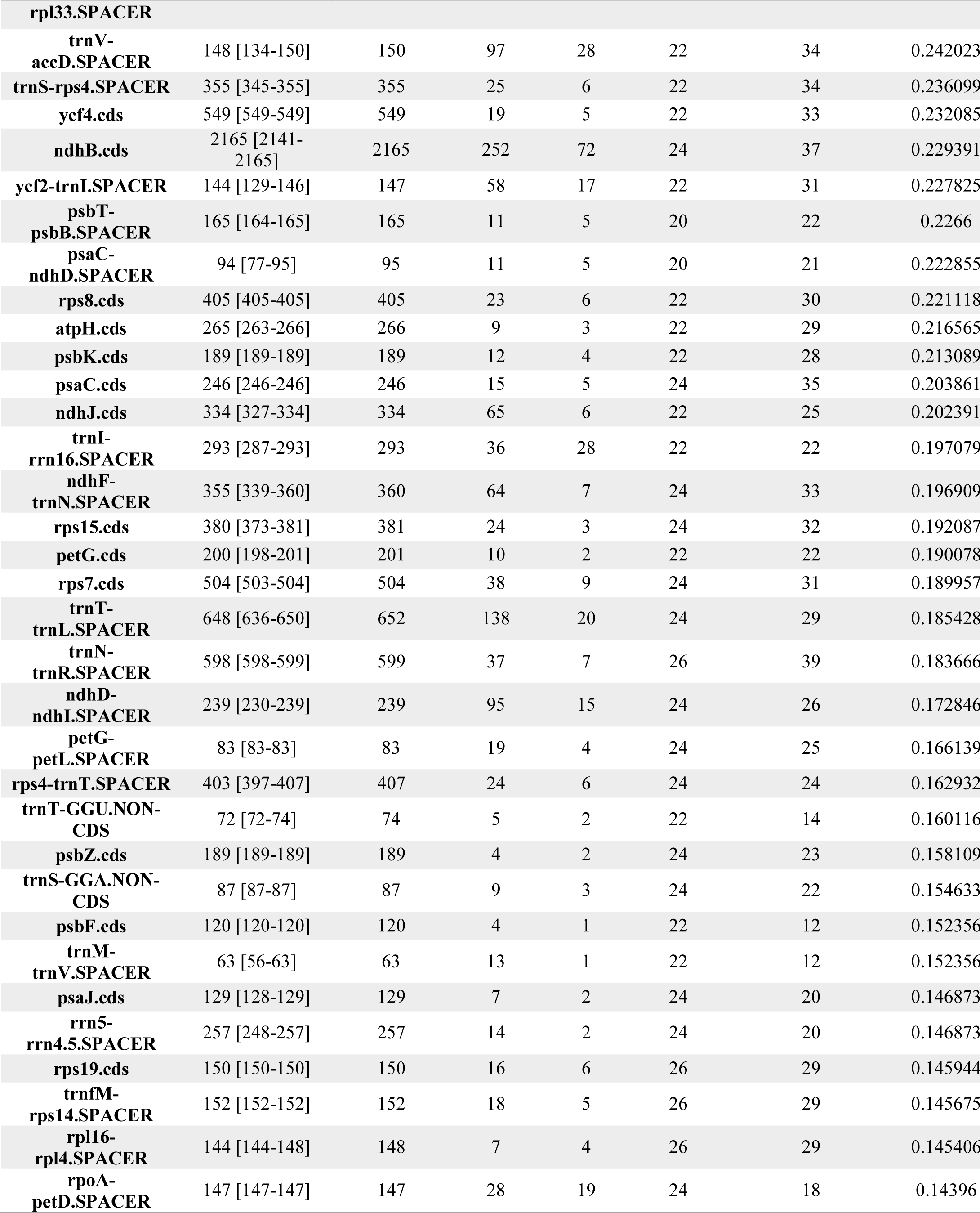
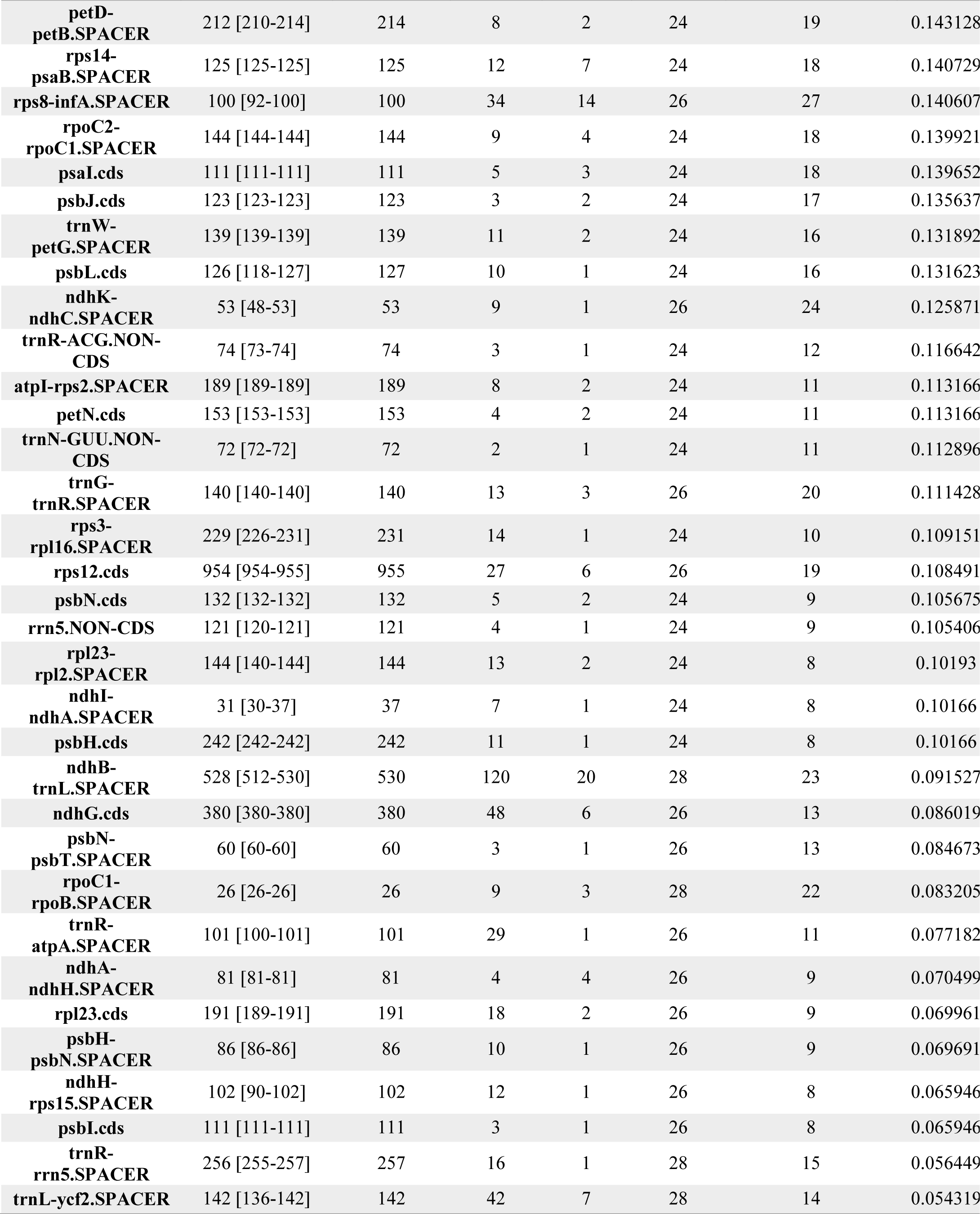
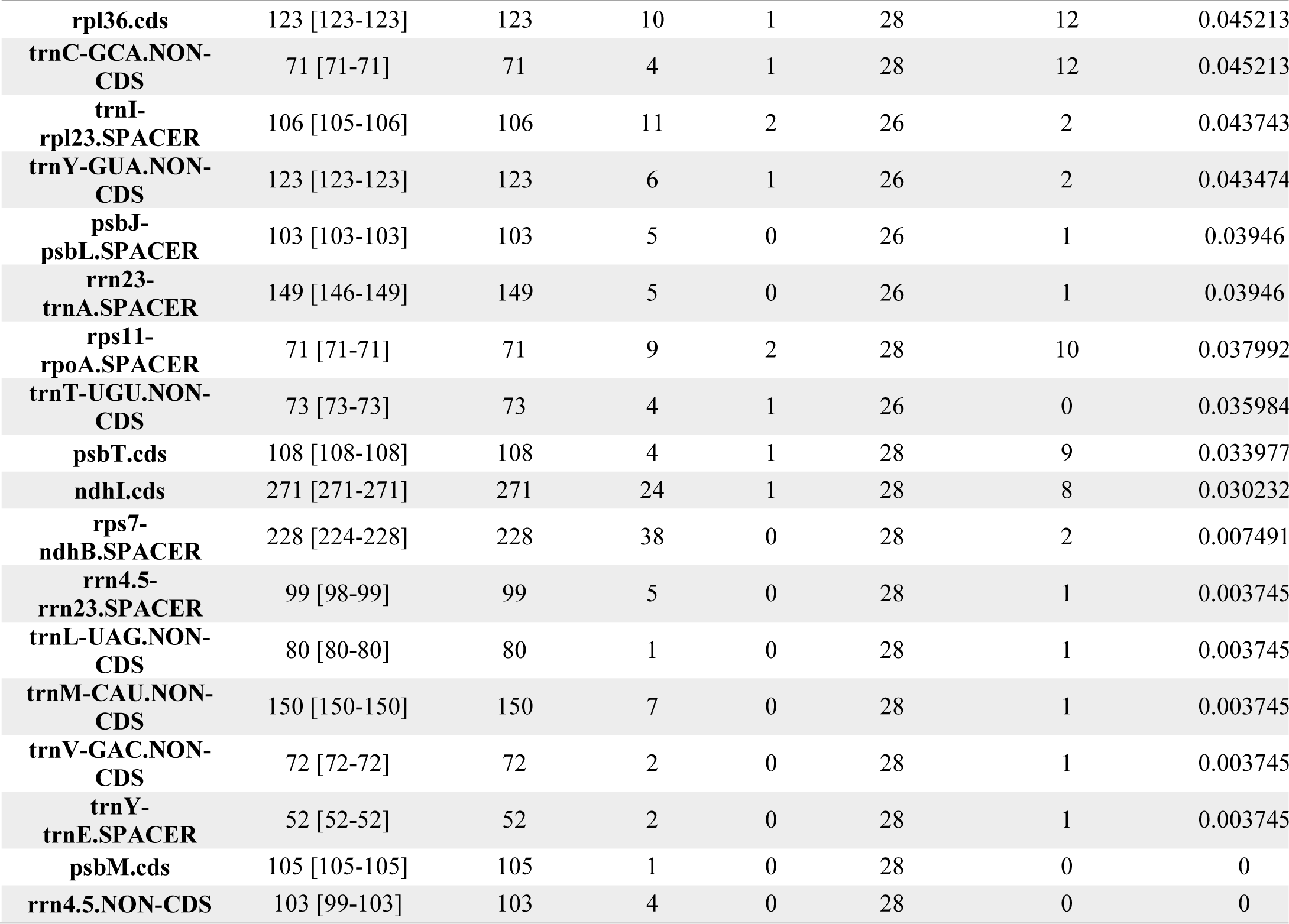
Summary statistics for all chloroplast markers. Markers are ranked by phylogenetic information based on a weighed mean of relative values of number of variable sites (weight = 1), mean bootstrap (weight = 2) and distance to the full plastid tree (weight = 3).

**Table S4.** Summary statistics for the top 10 markers with suitable size for PCR amplification. Markers are ranked by phylogenetic information based on a weighed mean of relative values of number of variable sites (weight = 1), mean bootstrap (weight = 2) and distance to the full plastid tree (weight = 3).

|  | **Bp** | **Aligned_bp** | **Variable** | **PIS** | **Distance** | **Bootstrap..Mean.** |
| --- | --- | --- | --- | --- | --- | --- |
| **clpP.cds** | 359 [356-362] | 362 | 112 | 64 | 8 | 70 |
| **ycf4-cemA.SPACER** | 752 [736-755] | 757 | 72 | 27 | 8 | 67 |
| **rps2.cds** | 729 [726-729] | 729 | 77 | 24 | 10 | 71 |
| **rbcL-atpB.SPACER** | 765 [746-768] | 768 | 122 | 27 | 12 | 79 |
| **psbB-clpP.SPACER** | 547 [525-553] | 556 | 100 | 41 | 10 | 67 |
| **petA.cds** | 963 [963-963] | 963 | 50 | 18 | 8 | 59 |
| **ycf3-trnS.SPACER** | 804 [790-805] | 805 | 67 | 22 | 14 | 82 |
| **rps3.cds** | 657 [657-657] | 657 | 47 | 16 | 10 | 63 |
| **ndhC-rbcL.SPACER** | 716 [705-717] | 718 | 103 | 13 | 8 | 49 |
| **ndhE-psaC.SPACER** | 318 [315-328] | 328 | 70 | 26 | 14 | 75 |

## References

Anderson, E.F. (2001). The cactus family. Portland: Timber Press.

APG (2016). An update of the Angiosperm Phylogeny Group classification for the orders and families of flowering plants: APG IV. Botanical Journal of the Linnean Society 181, 1–20. doi:10.1111/boj.12385.

Arakaki, M., Christin, P.-A., Nyffeler, R., Lendel, A., Eggli, U., Ogburn, R. M., et al. (2011). Contemporaneous and recent radiations of the world’s major succulent plant lineages. PNAS 108, 8379–8384. doi:10.1073/pnas.1100628108.

Backeberg, C. (1958). Die Cactaceae. Jena: Veb Gustav Fischer Verlag.

Bárcenas, R. T. (2016). A molecular phylogenetic approach to the systematics of Cylindropuntieae (Opuntioideae, Cactaceae). Cladistics 32, 351–359. doi:10.1111/cla.12135.

Bárcenas, R. T., Yesson, C., and Hawkins, J. A. (2011). Molecular systematics of the Cactaceae. Cladistics 27, 470–489. doi:10.1111/j.1096-0031.2011.00350.x.

Barfuss, M. H. J., Samuel, R., Till, W., and Stuessy, T. F. (2005). Phylogenetic relationships in subfamily Tillandsioideae (Bromeliaceae) based on DNA sequence data from seven plastid regions. Am. J. Bot. 92, 337–351. doi:10.3732/ajb.92.2.337.

Barres, L., Vilatersana, R., Molero, J., Susanna, A., and Galbany-Casals, M. (2011). Molecular phylogeny of Euphorbia subg. Esula sect. Aphyllis (Euphorbiaceae) inferred from nrDNA and cpDNA markers with biogeographic insights. Taxon 60, 705–720.

Bedoya, A. M., Ruhfel, B. R., Philbrick, C. T., Madriñán, S., Bove, C. P., Mesterházy, A., et al. (2019). Plastid Genomes of Five Species of Riverweeds (Podostemaceae): Structural Organization and Comparative Analysis in Malpighiales. Front. Plant Sci. 10. doi:10.3389/fpls.2019.01035.

Borsch, T., Korotkova, N., Raus, T., Lobin, W., and Löhne, C. (2009). The petD group II intron as a species level marker: utility for tree inference and species identification in the diverse genus Campanula (Campanulaceae). will 39, 7–33. doi:10.3372/wi.39.39101.

Bouillé, M., Senneville, S., and Bousquet, J. (2011). Discordant mtDNA and cpDNA phylogenies indicate geographic speciation and reticulation as driving factors for the diversification of the genus Picea. Tree Genetics & Genomes 7, 469–484. doi:10.1007/s11295-010-0349-z.

Braukmann, T. W. A., Kuzmina, M., and Stefanovi, S. (2009). Loss of all plastid ndh genes in Gnetales and conifers: extent and evolutionary significance for the seed plant phylogeny. Curr. Genet. 55, 323–337. doi:10.1007/s00294-009-0249-7.

Britton, N.L., and Rose, J.N. (1919). The Cactaceae, Volume I and Volume II. New York: Dover.

Brochmann, C., Brysting, A. K., Alsos, I. G., Borgen, L., Grundt, H. H., Scheen, A.-C., et al. (2004). Polyploidy in arctic plants. Biological Journal of the Linnean Society 82, 521–536. doi:10.1111/j.1095-8312.2004.00337.x.

Brudno, M., Malde, S., Poliakov, A., Do, C. B., Couronne, O., Dubchak, I., et al. (2003). Glocal alignment: finding rearrangements during alignment. Bioinformatics 19, i54–i62. doi:10.1093/bioinformatics/btg1005.

Bushnell, B. (2016). BBMap short read aligner, and other bioinformatic tools. https://sourceforge.net/projects/bbmap/

Cai, Z., Guisinger, M., Kim, H.-G., Ruck, E., Blazier, J. C., McMurtry, V., et al. (2008). Extensive reorganization of the plastid genome of Trifolium subterraneum (Fabaceae) is associated with numerous repeated sequences and novel DNA insertions. J. Mol. Evol. 67, 696–704. doi:10.1007/s00239-008-9180-7.

Carbonell-Caballero, J., Alonso, R., Ibañez, V., Terol, J., Talon, M., and Dopazo, J. (2015). A Phylogenetic Analysis of 34 Chloroplast Genomes Elucidates the Relationships between Wild and Domestic Species within the Genus Citrus. Mol Biol Evol 32, 2015–2035. doi:10.1093/molbev/msv082.

Cauz-Santos, L. A., Munhoz, C. F., Rodde, N., Cauet, S., Santos, A. A., Penha, H. A., et al. (2017). The Chloroplast Genome of Passiflora edulis (Passifloraceae) Assembled from Long Sequence Reads: Structural Organization and Phylogenomic Studies in Malpighiales. Front. Plant Sci. 8. doi:10.3389/fpls.2017.00334.

Cho, W.-B., Choi, B.-H., Kim, J.-H., Lee, D.-H., and Lee, J.-H. (2018). Complete Plastome Sequencing Reveals an Extremely Diminished SSC Region in Hemiparasitic Pedicularis ishidoyana (Orobanchaceae). anbf 55, 171–183. doi:10.5735/085.055.0122.

Chromosomal evolution in higher plants *Wellcome Library*. Available at: http://wellcomelibrary.org/item/b18036594 [Accessed January 28, 2020].

Chumley, T. W., Palmer, J. D., Mower, J. P., Fourcade, H. M., Calie, P. J., Boore, J. L., et al. (2006). The complete chloroplast genome sequence of Pelargonium x hortorum: organization and evolution of the largest and most highly rearranged chloroplast genome of land plants. Mol. Biol. Evol. 23, 2175–2190. doi:10.1093/molbev/msl089.

Cui, Y., Zhou, J., Chen, X., Xu, Z., Wang, Y., Sun, W., et al. (2019). Complete chloroplast genome and comparative analysis of three Lycium (Solanaceae) species with medicinal and edible properties. Gene Reports 17, 100464. doi:10.1016/j.genrep.2019.100464.

Daniell, H., Lin, C.-S., Yu, M., and Chang, W.-J. (2016). Chloroplast genomes: diversity, evolution, and applications in genetic engineering. Genome Biology 17, 134. doi:10.1186/s13059-016-1004-2.

Darling, A. C. E., Mau, B., Blattner, F. R., and Perna, N. T. (2004). Mauve: multiple alignment of conserved genomic sequence with rearrangements. Genome Res. 14, 1394–1403. doi:10.1101/gr.2289704.

Dastpak, A., Osaloo, S. K., Maassoumi, A. A., and Safar, K. N. (2018). Molecular Phylogeny of Astragalus sect. Ammodendron (Fabaceae) Inferred from Chloroplast ycf1 Gene. anbf 55, 75–82. doi:10.5735/085.055.0108.

Demaio, P. H., Barfuss, M. H. J., Kiesling, R., Till, W., and Chiapella, J. O. (2011). Molecular phylogeny of Gymnocalycium (Cactaceae): assessment of alternative infrageneric systems, a new subgenus, and trends in the evolution of the genus. Am. J. Bot. 98, 1841–1854. doi:10.3732/ajb.1100054.

Dierckxsens, N., Mardulyn, P., and Smits, G. (2017). NOVOPlasty: de novo assembly of organelle genomes from whole genome data. Nucleic Acids Res 45, e18–e18. doi:10.1093/nar/gkw955.

Dong, W., Liu, J., Yu, J., Wang, L., and Zhou, S. (2012). Highly Variable Chloroplast Markers for Evaluating Plant Phylogeny at Low Taxonomic Levels and for DNA Barcoding. PLOS ONE 7, e35071. doi:10.1371/journal.pone.0035071.

Dong, W., Xu, C., Cheng, T., and Zhou, S. (2013). Complete Chloroplast Genome of Sedum sarmentosum and Chloroplast Genome Evolution in Saxifragales. PLOS ONE 8, e77965. doi:10.1371/journal.pone.0077965.

Dong, W., Xu, C., Li, C., Sun, J., Zuo, Y., Shi, S., et al. (2015). ycf1, the most promising plastid DNA barcode of land plants. Sci Rep 5, 1–5. doi:10.1038/srep08348.

Dong, W.-L., Wang, R.-N., Zhang, N.-Y., Fan, W.-B., Fang, M.-F., and Li, Z.-H. (2018). Molecular Evolution of Chloroplast Genomes of Orchid Species: Insights into Phylogenetic Relationship and Adaptive Evolution. International Journal of Molecular Sciences 19, 716. doi:10.3390/ijms19030716.

Doyle, J.J., and Doyle, J.L. (1987). A rapid DNA isolation procedure for small quantities of fresh leaf tissue. Phytochemical Bulletin 19, 11–15.

Doweld, A.B. (1999). Tribal taxonomy of Pereskioideae and Opuntioideae (Cactaceae). Succulents 1(2), 25–26.

Downie, S. R., and Jansen, R. K. (2015). A Comparative Analysis of Whole Plastid Genomes from the Apiales: Expansion and Contraction of the Inverted Repeat, Mitochondrial to Plastid Transfer of DNA, and Identification of Highly Divergent Noncoding Regions. sbot 40, 336–351. doi:10.1600/036364415X686620.

Downie, S. R., Katz Downie, D. S., and Watson, M. F. (2000). A phylogeny of the flowering plant family Apiaceae based on chloroplast DNA rpl16 and rpoC1 intron sequences: towards a suprageneric classification of subfamily Apioideae. American Journal of Botany 87, 273–292. doi:10.2307/2656915.

Drescher, A., Ruf, S., Calsa, T., Carrer, H., and Bock, R. (2000). The two largest chloroplast genome-encoded open reading frames of higher plants are essential genes. Plant J. 22, 97–104. doi:10.1046/j.1365-313x.2000.00722.x.

Edwards, E. J., Nyffeler, R., and Donoghue, M. J. (2005). Basal cactus phylogeny: implications of Pereskia (Cactaceae) paraphyly for the transition to the cactus life form. Am. J. Bot. 92, 1177–1188. doi:10.3732/ajb.92.7.1177.

Ekenäs, C., Baldwin, B. G., and Andreasen, K. (2007). A Molecular Phylogenetic Study of Arnica (Asteraceae): Low Chloroplast DNA Variation and Problematic Subgeneric Classification. sbot 32, 917–928. doi:10.1043/06-80.1.

Fajardo, D., Senalik, D., Ames, M., Zhu, H., Steffan, S. A., Harbut, R., et al. (2013). Complete plastid genome sequence of Vaccinium macrocarpon: structure, gene content, and rearrangements revealed by next generation sequencing. Tree Genetics & Genomes 9, 489–498. doi:10.1007/s11295-012-0573-9.

Fonseca, L. H. M., and Lohmann, L. G. (2017). Plastome Rearrangements in the “Adenocalymma-Neojobertia” Clade (Bignonieae, Bignoniaceae) and Its Phylogenetic Implications. Front. Plant Sci. 8. doi:10.3389/fpls.2017.01875.

Frazer, K. A., Pachter, L., Poliakov, A., Rubin, E. M., and Dubchak, I. (2004). VISTA: computational tools for comparative genomics. Nucleic Acids Res. 32, W273–279. doi:10.1093/nar/gkh458.

Gardner, A. G., Sessa, E. B., Michener, P., Johnson, E., Shepherd, K. A., Howarth, D. G., et al. (2016). Utilizing next-generation sequencing to resolve the backbone of the Core Goodeniaceae and inform future taxonomic and floral form studies. Mol. Phylogenet. Evol. 94, 605–617. doi:10.1016/j.ympev.2015.10.003.

Gernandt, D. S., Hernández-León, S., Salgado-Hernández, E., and Rosa, J. A. P. de la (2009). Phylogenetic Relationships of Pinus Subsection Ponderosae Inferred from Rapidly Evolving cpDNA Regions. sbot 34, 481–491. doi:10.1600/036364409789271290.

Gitzendanner, M. A., Soltis, P. S., Yi, T.-S., Li, D.-Z., and Soltis, D. E. (2018). “Plastome Phylogenetics: 30 Years of Inferences Into Plant Evolution,” in Advances in Botanical Research Plastid Genome Evolution., eds. S.-M. Chaw and R. K. Jansen (Academic Press), 293–313. doi:10.1016/bs.abr.2017.11.016.

Gonçalves, D. J. P., Simpson, B. B., Ortiz, E. M., Shimizu, G. H., and Jansen, R. K. (2019). Incongruence between gene trees and species trees and phylogenetic signal variation in plastid genes. Mol. Phylogenet. Evol. 138, 219–232. doi:10.1016/j.ympev.2019.05.022.

GPWS (2001). Phylogeny and Subfamilial Classification of the Grasses (Poaceae). Annals of the Missouri Botanical Garden 88, 373–457. doi:10.2307/3298585.

Graham, S. W., Lam, V. K. Y., and Merckx, V. S. F. T. (2017). Plastomes on the edge: the evolutionary breakdown of mycoheterotroph plastid genomes. New Phytologist 214, 48–55. doi:10.1111/nph.14398.

Greiner, S., Lehwark, P., and Bock, R. (2019). OrganellarGenomeDRAW (OGDRAW) version 1.3.1: expanded toolkit for the graphical visualization of organellar genomes. Nucleic Acids Res. 47, W59–W64. doi:10.1093/nar/gkz238.

Griffith, M. P., and Porter, J. M. (2009). Phylogeny of Opuntioideae (Cactaceae). International Journal of Plant Sciences 170, 107–116. doi:10.1086/593048.

Guerrero, P. C., Majure, L. C., Cornejo-Romero, A., and Hernández-Hernández, T. (2019). Phylogenetic Relationships and Evolutionary Trends in the Cactus Family. J Hered 110, 4–21. doi:10.1093/jhered/esy064.

Guo, J., and Wilson, C. A. (2013). Molecular Phylogeny of Crested Iris Based on Five Plastid Markers (Iridaceae). Sbot 38, 987–995. doi:10.1600/036364413X674724.

Guo, Y.-Y., Luo, Y.-B., Liu, Z.-J., and Wang, X.-Q. (2012). Evolution and Biogeography of the Slipper Orchids: Eocene Vicariance of the Conduplicate Genera in the Old and New World Tropics. PLOS ONE 7, e38788. doi:10.1371/journal.pone.0038788.

Haberle, R. C., Fourcade, H. M., Boore, J. L., and Jansen, R. K. (2008). Extensive Rearrangements in the Chloroplast Genome of Trachelium caeruleum Are Associated with Repeats and tRNA Genes. J Mol Evol 66, 350–361. doi:10.1007/s00239-008-9086-4.

Harris, M. E., Meyer, G., Vandergon, T., and Vandergon, V. O. (2013). Loss of the Acetyl-CoA Carboxylase (accD) Gene in Poales. Plant Mol Biol Rep 31, 21–31. doi:10.1007/s11105-012-0461-3.

He, Y., Hou, P., Fan, G., Arain, S., and Peng, C. (2014). Comprehensive analyses of molecular phylogeny and main alkaloids for Coptis (Ranunculaceae) species identification. Biochemical Systematics and Ecology 56, 88–94. doi:10.1016/j.bse.2014.05.002.

Hernández D Hernández, T., Brown, J. W., Schlumpberger, B. O., Eguiarte, L. E., and Magallón, S. (2014). Beyond aridification: multiple explanations for the elevated diversification of cacti in the New World Succulent Biome. New Phytologist 202, 1382–1397. doi:10.1111/nph.12752.

Hernández-Hernández, T., Hernández, H. M., De-Nova, J. A., Puente, R., Eguiarte, L. E., and Magallón, S. (2011). Phylogenetic relationships and evolution of growth form in Cactaceae (Caryophyllales, Eudicotyledoneae). Am. J. Bot. 98, 44–61. doi:10.3732/ajb.1000129.

Hu, S., Sablok, G., Wang, B., Qu, D., Barbaro, E., Viola, R., et al. (2015). Plastome organization and evolution of chloroplast genes in Cardamine species adapted to contrasting habitats. BMC Genomics 16, 306. doi:10.1186/s12864-015-1498-0.

Huang, S., Chiang, Y. C., Schaal, B. A., Chou, C. H., and Chiang, T. Y. (2001). Organelle DNA phylogeography of Cycas taitungensis, a relict species in Taiwan. Mol. Ecol. 10, 2669–2681. doi:10.1046/j.0962-1083.2001.01395.x.

Hunt, D. (2002). That’s *Opuntia*, that was! *Succul*. Plant Res. 6: 245–249.

Hunt, D., Taylor, N., and Charles, G. (2006). The New Cactus Lexicon. Milborne Port: DH Books.

Hunt, D. (2011). Classification of the ‘cylindroid’ opuntias of South America. Cactaceae Systematics Initiatives 25, 5–29.

Iles, W. J. D., Smith, S. Y., and Graham, S. W. (2013). A well-supported phylogenetic framework for the monocot order Alismatales reveals multiple losses of the plastid NADH dehydrogenase complex and a strong long-branch effect. Early Events in Monocot Evolution. Cambridge: Cambridge University Press. doi:10.1017/CBO9781139002950.002.

Jansen, R. K., and Ruhlman, T. A. (2012). “Plastid Genomes of Seed Plants,” in Genomics of Chloroplasts and Mitochondria Advances in Photosynthesis and Respiration., eds. R. Bock and V. Knoop (Dordrecht: Springer Netherlands), 103–126. doi:10.1007/978-94-007-2920-9_5.

Jansen, R. K., Saski, C., Lee, S.-B., Hansen, A. K., and Daniell, H. (2011). Complete plastid genome sequences of three Rosids (Castanea, Prunus, Theobroma): evidence for at least two independent transfers of rpl22 to the nucleus. Mol. Biol. Evol. 28, 835–847. doi:10.1093/molbev/msq261.

Jaramillo, M. A., Callejas, R., Davidson, C., Smith, J. F., Stevens, A. C., and Tepe, E. J. (2008). A Phylogeny of the Tropical Genus Piper Using Its and the Chloroplast Intron psbJ–petA. sbot 33, 647–660. doi:10.1600/036364408786500244.

Jin, D.-P., Choi, I.-S., and Choi, B.-H. (2019). Plastid genome evolution in tribe Desmodieae (Fabaceae: Papilionoideae). PLOS ONE 14, e0218743. doi:10.1371/journal.pone.0218743.

Katoh, K., and Standley, D. M. (2013). MAFFT multiple sequence alignment software version 7: improvements in performance and usability. Mol. Biol. Evol. 30, 772–780. doi:10.1093/molbev/mst010.

Kikuchi, S., Bédard, J., Hirano, M., Hirabayashi, Y., Oishi, M., Imai, M., et al. (2013). Uncovering the protein translocon at the chloroplast inner envelope membrane. Science 339, 571–574. doi:10.1126/science.1229262.

Kim, D.-K., Kim, J. S., and Kim, J.-H. (2012). The phylogenetic relationships of Asparagales in Korea based on five plastid DNA regions. J. Plant Biol. 55, 325–341. doi:10.1007/s12374-011-0016-4.

Kim, H. T., Chung, M. G., and Kim, K.-J. (2014). Chloroplast genome evolution in early diverged leptosporangiate ferns. Mol. Cells 37, 372–382. doi:10.14348/molcells.2014.2296.

Kim, H. T., Kim, J. S., Moore, M. J., Neubig, K. M., Williams, N. H., Whitten, W. M., et al. (2015). Seven New Complete Plastome Sequences Reveal Rampant Independent Loss of the ndh Gene Family across Orchids and Associated Instability of the Inverted Repeat/Small Single-Copy Region Boundaries. PLOS ONE 10, e0142215. doi:10.1371/journal.pone.0142215.

Kode, V., Mudd, E. A., Iamtham, S., and Day, A. (2005). The tobacco plastid accD gene is essential and is required for leaf development. The Plant Journal 44, 237–244. doi:10.1111/j.1365-313X.2005.02533.x.

Korotkova, N., Zabel, L., Quandt, D., and Barthlott, W. (2010). A phylogenetic analysis of Pfeiffera and the reinstatement of Lymanbensonia as an independently evolved lineage of epiphytic Cactaceae within a new tribe Lymanbensonieae. will 40, 151–172. doi:10.3372/wi.40.40201.

Kulshreshtha, S., Creamer, R., and Sterling, T. M. (2004). Phylogenetic relationships among New Mexico Astragalus mollissimus varieties and Oxytropis species by restriction fragment analysis. Weed Science 52, 984–988. doi:10.1614/WS-03-143R1.

Kuroda, H., and Maliga, P. (2003). The plastid clpP1 protease gene is essential for plant development. Nature 425, 86–89. doi:10.1038/nature01909.

Lam, V. K. Y., Merckx, V. S. F. T., and Graham, S. W. (2016). A few-gene plastid phylogenetic framework for mycoheterotrophic monocots. American Journal of Botany 103, 692–708. doi:10.3732/ajb.1500412.

Landrum, J. V. (2002). Four Succulent Families and 40 Million Years of Evolution and Adaptation to Xeric Environments: What Can Stem and Leaf Anatomical Characters Tell Us about Their Phylogeny? Taxon 51, 463– 473. doi:10.2307/1554859.

Las Peñas, M. L., Kiesling, R., and Bernardello, G. (2019). Phylogenetic reconstruction of the genus Tephrocactus (Cactaceae) based on molecular, morphological, and cytogenetical data. TAXON 68, 714–730. doi:10.1002/tax.12092.

Laslett, D., and Canback, B. (2004). ARAGORN, a program to detect tRNA genes and tmRNA genes in nucleotide sequences. Nucleic Acids Res. 32, 11–16. doi:10.1093/nar/gkh152.

Li, Y., Zhou, J., Chen, X., Cui, Y., Xu, Z., Li, Y., et al. (2017). Gene losses and partial deletion of small single-copy regions of the chloroplast genomes of two hemiparasitic Tax i llus species. Sci Rep 7, 1–12. doi:10.1038/s41598-017-13401-4.

Liston, A., and Wheeler, J. A. (1994). The phylogenetic position of the genus Astragalus (fabaceae): Evidence from the chloroplast genes rpoC1 and rpoC2. Biochemical Systematics and Ecology 22, 377–388. doi:10.1016/0305-1978(94)90028-0.

Liu, T.-J., Zhang, C.-Y., Yan, H.-F., Zhang, L., Ge, X.-J., and Hao, G. (2016). Complete plastid genome sequence of Primula sinensis (Primulaceae): structure comparison, sequence variation and evidence for accD transfer to nucleus. PeerJ 4, e2101. doi:10.7717/peerj.2101.

Liu, X., Yang, H., Zhao, J., Zhou, B., Li, T., and Xiang, B. (2017). The complete chloroplast genome sequence of the folk medicinal and vegetable plant purslane (Portulaca oleracea L.). The Journal of Horticultural Science and Biotechnology 93, 356–365. doi:10.1080/14620316.2017.1389308.

Loera, I., Sosa, V., and Ickert-Bond, S. M. (2012). Diversification in North American arid lands: Niche conservatism, divergence and expansion of habitat explain speciation in the genus Ephedra. Molecular Phylogenetics and Evolution 65, 437–450. doi:10.1016/j.ympev.2012.06.025.

Logacheva, M. D., Penin, A. A., Samigullin, T. H., Vallejo-Roman, C. M., and Antonov, A. S. (2007). Phylogeny of flowering plants by the chloroplast genome sequences: in search of a “lucky gene.” Biochemistry Moscow 72, 1324– 1330. doi:10.1134/S0006297907120061.

Logacheva, M. D., Schelkunov, M. I., Nuraliev, M. S., Samigullin, T. H., and Penin, A. A. (2014). The Plastid Genome of Mycoheterotrophic Monocot Petrosavia stellaris Exhibits Both Gene Losses and Multiple Rearrangements. Genome Biol Evol 6, 238–246. doi:10.1093/gbe/evu001.

Löhne, C., Borsch, T., and Wiersema, J. H. (2007). Phylogenetic analysis of Nymphaeales using fast-evolving and noncoding chloroplast markers. Bot J Linn Soc 154, 141–163. doi:10.1111/j.1095-8339.2007.00659.x.

Lohse, M., Drechsel, O., Kahlau, S., and Bock, R. (2013). OrganellarGenomeDRAW--a suite of tools for generating physical maps of plastid and mitochondrial genomes and visualizing expression data sets. Nucleic Acids Res. 41, W575–581. doi:10.1093/nar/gkt289.

Lowe, T. M., and Chan, P. P. (2016). tRNAscan-SE On-line: integrating search and context for analysis of transfer RNA genes. Nucleic Acids Res. 44, W54–57. doi:10.1093/nar/gkw413.

Lu, R.-S., Li, P., and Qiu, Y.-X. (2017). The Complete Chloroplast Genomes of Three Cardiocrinum (Liliaceae) Species: Comparative Genomic and Phylogenetic Analyses. Front. Plant Sci. 7. doi:10.3389/fpls.2016.02054.

Ma, P.-F., Zhang, Y.-X., Zeng, C.-X., Guo, Z.-H., and Li, D.-Z. (2014). Chloroplast Phylogenomic Analyses Resolve Deep-Level Relationships of an Intractable Bamboo Tribe Arundinarieae (Poaceae). Syst Biol 63, 933–950. doi:10.1093/sysbio/syu054.

Machado, L. de O., Vieira, L. do N., Stefenon, V. M., Oliveira Pedrosa, F. de, Souza, E. M. de, Guerra, M. P., et al. (2017). Phylogenomic relationship of feijoa (Acca sellowiana (O.Berg) Burret) with other Myrtaceae based on complete chloroplast genome sequences. Genetica 145, 163–174. doi:10.1007/s10709-017-9954-1.

Magee, A. M., Aspinall, S., Rice, D. W., Cusack, B. P., Sémon, M., Perry, A. S., et al. (2010). Localized hypermutation and associated gene losses in legume chloroplast genomes. Genome Res. 20, 1700–1710. doi:10.1101/gr.111955.110.

Majure, L.C., Baker, M. A., Cloud-Hughes, M., Salywon, A., and Neubig, K. M. (2019a). Phylogenomics in Cactaceae: A case study using the chollas sensu lato (Cylindropuntieae, Opuntioideae) reveals a common pattern out of the Chihuahuan and Sonoran deserts. Am. J. Bot. 106, 1327–1345. doi:10.1002/ajb2.1364.

Majure, L.C., Baker, M. A., Cloud D Hughes, M., Salywon, A., and Neubig, K. M. (2019b). Phylogenomics in Cactaceae: A case study using the chollas sensu lato (Cylindropuntieae, Opuntioideae) reveals a common pattern out of the Chihuahuan and Sonoran deserts. American Journal of Botany 106, 1327–1345. doi:10.1002/ajb2.1364.

Majure, L.C., Puente, R., Griffith, M. P., Judd, W. S., Soltis, P. S., and Soltis, D. E. (2012). Phylogeny of Opuntia s.s. (Cactaceae): clade delineation, geographic origins, and reticulate evolution. Am. J. Bot. 99, 847–864. doi:10.3732/ajb.1100375.

Majure, L.C., and Puente, R. (2014). Phylogenetic relationships and morphological evolution in *Opuntia* s. str. and closely related members of tribe Opuntieae. Succul. Plant Res. 8: 9–30.

Manen, J.-F., Habashi, C., Jeanmonod, D., Park, J.-M., and Schneeweiss, G. M. (2004). Phylogeny and intraspecific variability of holoparasitic Orobanche (Orobanchaceae) inferred from plastid rbcL sequences. Molecular Phylogenetics and Evolution 33, 482–500. doi:10.1016/j.ympev.2004.06.010.

Marshall, H. D., Newton, C., and Ritland, K. (2001). Sequence-repeat polymorphisms exhibit the signature of recombination in lodgepole pine chloroplast DNA. Mol. Biol. Evol. 18, 2136–2138. doi:10.1093/oxfordjournals.molbev.a003757.

Marx, H. E., O’Leary, N., Yuan, Y.-W., Lu D Irving, P., Tank, D. C., Múlgura, M. E., et al. (2010). A molecular phylogeny and classification of Verbenaceae. American Journal of Botany 97, 1647–1663. doi:10.3732/ajb.1000144.

Mauseth, J. D. (1999). Anatomical Adaptations to Xeric Conditions in Maihuenia (Cactaceae), a Relictual, Leaf-Bearing Cactus. J Plant Res 112, 307–315. doi:10.1007/PL00013886.

McNeal, J. R., Bennett, J. R., Wolfe, A. D., and Mathews, S. (2013). Phylogeny and origins of holoparasitism in Orobanchaceae. American Journal of Botany 100, 971–983. doi:10.3732/ajb.1200448.

Michelangeli, F. A., Guimaraes, P. J. F., Penneys, D. S., Almeda, F., and Kriebel, R. (2013). Phylogenetic relationships and distribution of New World Melastomeae (Melastomataceae). Bot J Linn Soc 171, 38–60. doi:10.1111/j.1095-8339.2012.01295.x.

Miikeda, O., Kita, K., Handa, T., and Yukawa, T. (2006). Phylogenetic relationships of Clematis (Ranunculaceae) based on chloroplast and nuclear DNA sequences. Bot J Linn Soc 152, 153–168. doi:10.1111/j.1095-8339.2006.00551.x.

Miller, M. A., Pfeiffer, W., and Schwartz, T. (2010). Creating the CIPRES Science Gateway for inference of large phylogenetic trees. in 2010 Gateway Computing Environments Workshop (GCE), 1–8. doi:10.1109/GCE.2010.5676129.

Montes Moreno, N., Sáez, L., Benedí, C., Susanna, A., and Garcia-Jacas, N. (2010). Generic delineation, phylogeny and subtribal affinities of Phagnalon and Aliella (Compositae, Gnaphalieae) based on nuclear and chloroplast sequences. TAXON 59, 1654–1670. doi:10.1002/tax.596002.

Mooney, H. A., Gulmon, S. L., and Weisser, P. J. (1977). Environmental Adaptations of the Atacaman Desert Cactus Copiapoa haseltoniana. Flora 166, 117–124. doi:10.1016/S0367-2530(17)32124-2.

Morales D Briones, D. F., Arias, T., Stilio, V. S. D., and Tank, D. C. (2019). Chloroplast primers for clade-wide phylogenetic studies of Thalictrum. Applications in Plant Sciences 7, e11294. doi:10.1002/aps3.11294.

Mower, J. P., and Vickrey, T. L. (2018). “Structural Diversity Among Plastid Genomes of Land Plants,” in Advances in Botanical Research Plastid Genome Evolution., eds. S.-M. Chaw and R. K. Jansen (Academic Press), 263–292. doi:10.1016/bs.abr.2017.11.013.

Neubig, K. M., and Abbott, J. R. (2010). Primer development for the plastid region ycf1 in Annonaceae and other magnoliids. American Journal of Botany 97, e52–e55. doi:10.3732/ajb.1000128.

Neubig, K. M., Whitten, W. M., Carlsward, B. S., Blanco, M. A., Endara, L., Williams, N. H., et al. (2009). Phylogenetic utility of ycf1 in orchids: a plastid gene more variable than matK. Plant Syst Evol 277, 75–84. doi:10.1007/s00606-008-0105-0.

Neubig, K., Whitten, M., Abbott, R., Elliot, S., Soltis, D., Soltis, P., et al. (2014). Variables affecting DNA preservation in archival plant specimens. DNA Banking for the 21st Century.

Nobel, P.S. (2002). Cacti biology and uses. Berkley: University of California Press.

Nyffeler, R. (2002). Phylogenetic relationships in the cactus family (Cactaceae) based on evidence from trnK/ matK and trnL-trnF sequences. Am. J. Bot. 89, 312–326. doi:10.3732/ajb.89.2.312.

Oliver, M. J., Murdock, A. G., Mishler, B. D., Kuehl, J. V., Boore, J. L., Mandoli, D. F., et al. (2010). Chloroplast genome sequence of the moss Tortula ruralis: gene content, polymorphism, and structural arrangement relative to other green plant chloroplast genomes. BMC Genomics 11, 143. doi:10.1186/1471-2164-11-143.

Palmer, J. D., Jansen, R. K., Michaels, H. J., Chase, M. W., and Manhart, J. R. (1988). Chloroplast DNA Variation and Plant Phylogeny. Annals of the Missouri Botanical Garden 75, 1180–1206. doi:10.2307/2399279.

Panero, J. L., and Funk, V. A. (2002). Toward a phylogenetic subfamilial classification for the compositae (Asteraceae). PROC.BIOL.SOC.WASH. 115, 909–922.

Paradis, E., Claude, J., and Strimmer, K. (2004). APE: Analyses of Phylogenetics and Evolution in R language. Bioinformatics 20, 289–290. doi:10.1093/bioinformatics/btg412.

Park, I., Yang, S., Kim, W. J., Noh, P., Lee, H. O., and Moon, B. C. (2018). The Complete Chloroplast Genomes of Six Ipomoea Species and Indel Marker Development for the Discrimination of Authentic Pharbitidis Semen (Seeds of I. nil or I. purpurea). Front. Plant Sci. 9. doi:10.3389/fpls.2018.00965.

Park, S., Ruhlman, T. A., Weng, M.-L., Hajrah, N. H., Sabir, J. S. M., and Jansen, R. K. (2017). Contrasting Patterns of Nucleotide Substitution Rates Provide Insight into Dynamic Evolution of Plastid and Mitochondrial Genomes of Geranium. Genome Biol Evol 9, 1766–1780. doi:10.1093/gbe/evx124.

Parker, N., Wang, Y., and Meinke, D. (2014). Natural variation in sensitivity to a loss of chloroplast translation in Arabidopsis. Plant Physiol. 166, 2013–2027. doi:10.1104/pp.114.249052.

Peredo, E. L., King, U. M., and Les, D. H. (2013). The Plastid Genome of Najas flexilis: Adaptation to Submersed Environments Is Accompanied by the Complete Loss of the NDH Complex in an Aquatic Angiosperm. PLOS ONE 8, e68591. doi:10.1371/journal.pone.0068591.

Perry, A. S., and Wolfe, K. H. (2002). Nucleotide substitution rates in legume chloroplast DNA depend on the presence of the inverted repeat. J. Mol. Evol. 55, 501–508. doi:10.1007/s00239-002-2333-y.

Petersen, G., Seberg, O., Thorsøe, S., Jørgensen, T., and Mathew, B. (2008). A Phylogeny of the Genus Crocus (Iridaceae) Based on Sequence Data from Five Plastid Regions. Taxon 57, 487–499. doi:10.2307/25066017.

Peterson, P. M., Romaschenko, K., and Johnson, G. (2010). A classification of the Chloridoideae (Poaceae) based on multi-gene phylogenetic trees. Molecular Phylogenetics and Evolution 55, 580–598. doi:10.1016/j.ympev.2010.01.018.

Peterson, P. M., Romaschenko, K., Snow, N., and Johnson, G. (2012). A molecular phylogeny and classification of Leptochloa (Poaceae: Chloridoideae: Chlorideae) sensu lato and related genera. Ann Bot 109, 1317–1330. doi:10.1093/aob/mcs077.

Piot, A., Hackel, J., Christin, P.-A., and Besnard, G. (2018). One-third of the plastid genes evolved under positive selection in PACMAD grasses. Planta 247, 255–266. doi:10.1007/s00425-017-2781-x.

Prince, L. M. (2015). Plastid primers for angiosperm phylogenetics and phylogeography. Applications in Plant Sciences 3, 1400085. doi:10.3732/apps.1400085.

Raubeson, L. A., and Jansen, R. K. (2005). “Chloroplast genomes of plants.,” in Plant diversity and evolution: genotypic and phenotypic variation in higher plants, ed. R. J. Henry (Cambridge: University Press), 45–68. doi:10.1079/9780851999043.0045.

Reginato, M., Michelangeli, F. A., and Goldenberg, R. (2010). Phylogeny of Pleiochiton (Melastomataceae, Miconieae): total evidence. Botanical Journal of the Linnean Society 162, 423–434. doi:10.1111/j.1095-8339.2009.01022.x.

Reginato, M., Neubig, K. M., Majure, L. C., and Michelangeli, F. A. (2016). The first complete plastid genomes of Melastomataceae are highly structurally conserved. PeerJ 4, e2715. doi:10.7717/peerj.2715.

Ripma, L. A., Simpson, M. G., and Hasenstab-Lehman, K. (2014). Geneious! Simplified genome skimming methods for phylogenetic systematic studies: A case study in Oreocarya (Boraginaceae)1. Appl Plant Sci 2. doi:10.3732/apps.1400062.

Ritz, C. M., Reiker, J., Charles, G., Hoxey, P., Hunt, D., Lowry, M., et al. (2012). Molecular phylogeny and character evolution in terete-stemmed Andean opuntias (Cactaceae-Opuntioideae). Mol. Phylogenet. Evol. 65, 668–681. doi:10.1016/j.ympev.2012.07.027.

Robinson, D. F., and Foulds, L. R. (1981). Comparison of phylogenetic trees. Mathematical Biosciences 53, 131–147. doi:10.1016/0025-5564(81)90043-2.

Rousseau-Gueutin, M., Huang, X., Higginson, E., Ayliffe, M., Day, A., and Timmis, J. N. (2013). Potential functional replacement of the plastidic acetyl-CoA carboxylase subunit (accD) gene by recent transfers to the nucleus in some angiosperm lineages. Plant Physiol. 161, 1918–1929. doi:10.1104/pp.113.214528.

Rozas, J., Ferrer-Mata, A., Sánchez-DelBarrio, J. C., Guirao-Rico, S., Librado, P., Ramos-Onsins, S. E., et al. (2017). DnaSP 6: DNA Sequence Polymorphism Analysis of Large Data Sets. Mol. Biol. Evol. 34, 3299–3302. doi:10.1093/molbev/msx248.

Rozen, S., and Skaletsky, H. (2000). Primer3 on the WWW for general users and for biologist programmers. Methods Mol. Biol. 132, 365–386. doi:10.1385/1-59259-192-2:365.

Ruhlman, T. A., Chang, W.-J., Chen, J. J., Huang, Y.-T., Chan, M.-T., Zhang, J., et al. (2015). NDH expression marks major transitions in plant evolution and reveals coordinate intracellular gene loss. BMC Plant Biol 15, 100. doi:10.1186/s12870-015-0484-7.

Ruhlman, T. A., and Jansen, R. K. (2014). “The Plastid Genomes of Flowering Plants,” in Chloroplast Biotechnology: Methods and Protocols Methods in Molecular Biology., ed. P. Maliga (Totowa, NJ: Humana Press), 3–38. doi:10.1007/978-1-62703-995-6_1.

Ruhlman, T. A., and Jansen, R. K. (2018). “Aberration or Analogy? The Atypical Plastomes of Geraniaceae,” in Advances in Botanical Research Plastid Genome Evolution., eds. S.-M. Chaw and R. K. Jansen (Academic Press), 223–262. doi:10.1016/bs.abr.2017.11.017.

Sanderson, M. J., Copetti, D., Búrquez, A., Bustamante, E., Charboneau, J. L. M., Eguiarte, L. E., et al. (2015). Exceptional reduction of the plastid genome of saguaro cactus (Carnegiea gigantea): Loss of the ndh gene suite and inverted repeat. Am. J. Bot. 102, 1115–1127. doi:10.3732/ajb.1500184.

Särkinen, T., and George, M. (2013). Predicting Plastid Marker Variation: Can Complete Plastid Genomes from Closely Related Species Help? PLOS ONE 8, e82266. doi:10.1371/journal.pone.0082266.

Scataglini, M. A., Zuloaga, F. O., Giussani, L. M., Denham, S. S., and Morrone, O. (2014). Phylogeny of New World Paspalum (Poaceae, Panicoideae, Paspaleae) based on plastid and nuclear markers. Plant Syst Evol 300, 1051–1070. doi:10.1007/s00606-013-0944-1.

Schelkunov, M. I., Shtratnikova, V. Y., Nuraliev, M. S., Selosse, M.-A., Penin, A. A., and Logacheva, M. D. (2015). Exploring the limits for reduction of plastid genomes: a case study of the mycoheterotrophic orchids Epipogium aphyllum and Epipogium roseum. Genome Biol Evol 7, 1179–1191. doi:10.1093/gbe/evv019.

Schliep, K. P. (2011). phangorn: phylogenetic analysis in R. Bioinformatics 27, 592–593. doi:10.1093/bioinformatics/btq706.

Schumann, K. (1899). Gesambteschreibung der Kakteen. Neudamm: Verlag von J. Neumann. Shi, S., Li, J., Sun, J., Yu, J., and Zhou, S. (2013). Phylogeny and Classification of Prunus sensu lato (Rosaceae). Journal of Integrative Plant Biology 55, 1069–1079. doi:10.1111/jipb.12095.

Simmons, M. P. (2004). Independence of alignment and tree search. Molecular Phylogenetics and Evolution 31, 874–879. doi:10.1016/j.ympev.2003.10.008.

Solórzano, S., Chincoya, D. A., Sanchez-Flores, A., Estrada, K., Díaz-Velásquez, C. E., González-Rodríguez, A., et al. (2019). De Novo Assembly Discovered Novel Structures in Genome of Plastids and Revealed Divergent Inverted Repeats in Mammillaria (Cactaceae, Caryophyllales). Plants (Basel) 8. doi:10.3390/plants8100392.

Soltis, P. S., and Soltis, D. E. (2000). The role of genetic and genomic attributes in the success of polyploids. PNAS 97, 7051–7057. doi:10.1073/pnas.97.13.7051.

Stamatakis, A. (2014). RAxML version 8: a tool for phylogenetic analysis and post-analysis of large phylogenies. Bioinformatics 30, 1312–1313. doi:10.1093/bioinformatics/btu033.

Stefanović, S., Pfeil, B. E., Palmer, J. D., and Doyle, J. J. (2009). Relationships among Phaseoloid Legumes Based on Sequences from Eight Chloroplast Regions. Systematic Botany 34, 115–128.

Straub, S. C. K., Parks, M., Weitemier, K., Fishbein, M., Cronn, R. C., and Liston, A. (2012). Navigating the tip of the genomic iceberg: Next-generation sequencing for plant systematics. Am. J. Bot. 99, 349–364. doi:10.3732/ajb.1100335.

Stuppy, W. (2002). Seed characters and the generic classification of the Opuntioideae (Cactaceae). Succul. Plant Research 6, 25–58.

Sun, Y., Wang, A., Wan, D., Wang, Q., and Liu, J. (2012). Rapid radiation of Rheum (Polygonaceae) and parallel evolution of morphological traits. Molecular Phylogenetics and Evolution 63, 150–158. doi:10.1016/j.ympev.2012.01.002.

Taylor, N.P., Stuppy, W., and Barthlott, W. (2002). Realignment and revision of the Opuntioideae of Eastern Brazil. Succul. Plant Research 6, 99–132.

Thode, V. A., and Lohmann, L. G. (2019). Comparative Chloroplast Genomics at Low Taxonomic Levels: A Case Study Using Amphilophium (Bignonieae, Bignoniaceae). Front. Plant Sci. 10. doi:10.3389/fpls.2019.00796.

Thomson, A. M., Vargas, O. M., and Dick, C. W. (2018). Complete plastome sequences from Bertholletia excelsa and 23 related species yield informative markers for Lecythidaceae. Applications in Plant Sciences 6, e01151. doi:10.1002/aps3.1151.

Tillich, M., Lehwark, P., Pellizzer, T., Ulbricht-Jones, E. S., Fischer, A., Bock, R., et al. (2017). GeSeq - versatile and accurate annotation of organelle genomes. Nucleic Acids Res. 45, W6–W11. doi:10.1093/nar/gkx391.

Tsumura, Y., Kawahara, T., Wickneswari, R., and Yoshimura, K. (1996). Molecular phylogeny of Dipterocarpaceae in Southeast Asia using RFLP of PCR-amplified chloroplast genes. Theoret. Appl. Genetics 93, 22–29. doi:10.1007/BF00225722.

Vargas-Luna, M. D., Hernández-Ledesma, P., Majure, L. C., Puente-Martínez, R., Macías, H. M. H., and Bárcenas, R. T. (2018). Splitting Echinocactus: morphological and molecular evidence support the recognition of Homalocephala as a distinct genus in the Cacteae. PhytoKeys, 31–59. doi:10.3897/phytokeys.111.26856.

Vilatersana, R., Garcia-Jacas, N., Garnatje, T., Molero, J., Sonnante, G., and Susanna, A. (2010). Molecular Phylogeny of the Genus Ptilostemon (Compositae: Cardueae) and Its Relationships with Cynara and Lamyropsis. sbot 35, 907– 917. doi:10.1600/036364410X539952.

Wallace, R.S. (1995). Molecular systematic study of the Cactaceae: using chloroplast DNA variation to elucidate cactus phylogeny. Bradleya 13, 1–12. doi:https://doi.org/10.25223/brad.n13.1995.a1

Wallace, R.S., and Dickie, S.L. (2002). Systematic implication of chloroplast DNA sequence variation in subfamily Opuntioideae (Cactaceae). Succul. Plant Research 6, 9–24.

Walker, J. F., Jansen, R. K., Zanis, M. J., and Emery, N. C. (2015). Sources of inversion variation in the small single copy (SSC) region of chloroplast genomes. Am. J. Bot. 102, 1751–1752. doi:10.3732/ajb.1500299.

Walker, J. F., Walker-Hale, N., Vargas, O. M., Larson, D. A., and Stull, G. W. (2019). Characterizing gene tree conflict in plastome-inferred phylogenies. PeerJ 7, e7747. doi:10.7717/peerj.7747.

Walker, J. F., Yang, Y., Feng, T., Timoneda, A., Mikenas, J., Hutchison, V., et al. (2018). From cacti to carnivores: Improved phylotranscriptomic sampling and hierarchical homology inference provide further insight into the evolution of Caryophyllales. Am. J. Bot. 105, 446–462. doi:10.1002/ajb2.1069.

Wang, N., Yang, Y., Moore, M. J., Brockington, S. F., Walker, J. F., Brown, J. W., et al. (2019). Evolution of Portulacineae Marked by Gene Tree Conflict and Gene Family Expansion Associated with Adaptation to Harsh Environments. Mol Biol Evol 36, 112–126. doi:10.1093/molbev/msy200.

Wang, W., Li, H.-L., Xiang, X.-G., and Chen, Z.-D. (2014). Revisiting the phylogeny of Ranunculeae: Implications for divergence time estimation and historical biogeography. Journal of Systematics and Evolution 52, 551–565. doi:10.1111/jse.12101.

Weng, M.-L., Blazier, J. C., Govindu, M., and Jansen, R. K. (2014). Reconstruction of the ancestral plastid genome in Geraniaceae reveals a correlation between genome rearrangements, repeats, and nucleotide substitution rates. Mol. Biol. Evol. 31, 645–659. doi:10.1093/molbev/mst257.

Weng, M.-L., Ruhlman, T. A., and Jansen, R. K. (2016). Plastid-Nuclear Interaction and Accelerated Coevolution in Plastid Ribosomal Genes in Geraniaceae. Genome Biol Evol 8, 1824–1838. doi:10.1093/gbe/evw115.

Weng, M.-L., Ruhlman, T. A., and Jansen, R. K. (2017). Expansion of inverted repeat does not decrease substitution rates in Pelargonium plastid genomes. New Phytologist 214, 842–851. doi:10.1111/nph.14375.

Whitten, W. M., Neubig, K. M., and Williams, N. H. (2013). Generic and Subtribal relationShipS in neotropical cymbidieae (orchidaceae) baSed on matK/ycf1 plaStid data. 1. doi:10.15517/lank.v13i3.14425.

Wicke, S., Schneeweiss, G. M., dePamphilis, C. W., Müller, K. F., and Quandt, D. (2011). The evolution of the plastid chromosome in land plants: gene content, gene order, gene function. Plant Mol Biol 76, 273–297. doi:10.1007/s11103-011-9762-4.

Wommack, K. E., Bhavsar, J., and Ravel, J. (2008). Metagenomics: Read Length Matters. Appl Environ Microbiol 74, 1453–1463. doi:10.1128/AEM.02181-07.

Worberg, A., Quandt, D., Barniske, A.-M., Löhne, C., Hilu, K. W., and Borsch, T. (2007). Phylogeny of basal eudicots: Insights from non-coding and rapidly evolving DNA. Organisms Diversity & Evolution 7, 55–77. doi:10.1016/j.ode.2006.08.001.

Wu, F.-H., Chan, M.-T., Liao, D.-C., Hsu, C.-T., Lee, Y.-W., Daniell, H., et al. (2010). Complete chloroplast genome of Oncidium Gower Ramsey and evaluation of molecular markers for identification and breeding in Oncidiinae. BMC Plant Biology 10, 68. doi:10.1186/1471-2229-10-68.

Xi, Z., Ruhfel, B. R., Schaefer, H., Amorim, A. M., Sugumaran, M., Wurdack, K. J., et al. (2012). Phylogenomics and a posteriori data partitioning resolve the Cretaceous angiosperm radiation Malpighiales. PNAS 109, 17519–17524. doi:10.1073/pnas.1205818109.

Yao, G., Jin, J.-J., Li, H.-T., Yang, J.-B., Mandala, V. S., Croley, M., et al. (2019). Plastid phylogenomic insights into the evolution of Caryophyllales. Molecular Phylogenetics and Evolution 134, 74–86. doi:10.1016/j.ympev.2018.12.023.

Taylor, N., and Zappi, D. (2004). Cacti of Eastern Brazil. London: Royal Botanic Gardens, Kew.

Zhang, S., Soltis, D. E., Yang, Y., Li, D., and Yi, T. (2011). Multi-gene analysis provides a well-supported phylogeny of Rosales. Molecular Phylogenetics and Evolution 60, 21–28. doi:10.1016/j.ympev.2011.04.008.

Zhang, X.-Z., Zeng, C.-X., Ma, P.-F., Haevermans, T., Zhang, Y.-X., Zhang, L.-N., et al. (2016). Multi-locus plastid phylogenetic biogeography supports the Asian hypothesis of the temperate woody bamboos (Poaceae: Bambusoideae). Molecular Phylogenetics and Evolution 96, 118–129. doi:10.1016/j.ympev.2015.11.025.

Zong, D., Gan, P., Zhou, A., Zhang, Y., Zou, X., Duan, A., et al. (2019). Plastome Sequences Help to Resolve Deep-Level Relationships of Populus in the Family Salicaceae. Front. Plant Sci. 10. doi:10.3389/fpls.2019.00005.

